# Molecular characterization, localization, and physiological roles of ITP and ITP-L in the mosquito, *Aedes aegypti*

**DOI:** 10.1101/2024.01.21.576557

**Authors:** Farwa Sajadi, Jean-Paul Paluzzi

**Affiliations:** Department of Biology, York University, Toronto, Ontario, Canada

**Author notes:** **Correspondence**: Jean-Paul Paluzzi.

**Keywords:** ion transport peptide, ionoregulation, feeding, reproductive biology, anti-diuretic

## Abstract

The insect ion transport peptide (ITP) and its alternatively spliced variant, ITP-like peptide (ITP-L), belong to the crustacean hyperglycemic hormone family of peptides and are widely conserved among insect species. While limited, studies have characterized the ITP/ITP-L signaling system within insects, and putative functions including regulation of ion and fluid transport, ovarian maturation, and thirst/excretion have been proposed. Herein, we aimed to molecularly investigate *Itp* and *Itp-l* expression profiles in the mosquito *Aedes aegypti,* examine peptide immunolocalization and distribution within the adult central nervous system, and elucidate physiological roles for these neuropeptides. Transcript expression profiles of both *AedaeItp* and *AedaeItp-l* revealed distinct enrichment patterns in adults, with *AedaeItp* expressed in the brain and *AedaeItp-l* expression predominantly within the abdominal ganglia. Immunohistochemical analysis within the central nervous system revealed expression of *Aedae*ITP peptide in a number of cells in the brain and in the terminal ganglion. Comparatively, *Aedae*ITP-L peptide was localized solely within the pre-terminal abdominal ganglia of the central nervous system. Interestingly, prolonged desiccation stress caused upregulation of *AedaeItp* and *AedaeItp-l* levels in adult mosquitoes, suggesting possible functional roles in water conservation and feeding-related activities. RNAi-mediated knockdown of *AedaeItp* caused an increase in urine excretion, while knockdown of both *AedaeItp* and *AedaeItp-l* reduced blood feeding and egg-laying in females as well as hindered egg viability, suggesting roles in reproductive physiology and behaviour. Altogether, this study identifies *Aedae*ITP and *Aedae*ITP-L as key pleiotropic hormones, regulating various critical physiological processes in the disease vector, *A. aegypti*.

## Introduction

Neuropeptides comprise a large and diverse class of signaling molecules that, together with their receptors, play a significant role in controlling a myriad of behavioural and physiological processes, including reproduction, feeding, development, energy homeostasis, water and ion balance, and more (Coast, 2007; Schoofs et al., 2017; Strand et al., 2016) (Nässel et al., 2019). The crustacean hyperglycemic hormone (CHH) family of peptides are a large neuropeptide superfamily that includes structurally-related peptides composed of 72 to more than 80 amino acids (Chen et al., 2020) containing three highly conserved intramolecular disulfide bonds (Nagai et al., 2014). Functional roles for the CHH family of peptides are linked to molting, stress responses, reproduction, and homeostatic regulation of energy metabolism (Webster et al., 2012). The insect ion transport peptide (ITP) and its alternatively spliced variant, ITP-like or ITP-long (ITP-L) belong to the CHH family of peptides (Dai et al., 2007; Keller, 1992) and are widely conserved among insect species, including lepidopterans, such as the silkworm *Bombyx mori* (Dai et al., 2007; Endo et al., 2000), and the tobacco hornworm *Manduca sexta* (Drexler et al., 2007). While investigations examining the roles of ITP and ITP-L are limited, studies have suggested functions of both peptides as regulators of ion and fluid transport across the ileum of the desert locust *Schistocerca gregaria* (Audsley et al., 1992a), ecdysis in *M. sexta* (Drexler et al., 2007), ovarian maturation in the red flower beetle *Tribolium castaneum* (Begum et al., 2009), and thirst/excretion regulation and clock neuron modulation in the fruit fly *Drosophila melanogaster* (Gáliková et al., 2018; Johard et al., 2009).

The insect ITP was originally identified in *S. gregaria* (Audsley et al., 1992a; Audsley et al., 1992b; Dai et al., 2007), where it drives chloride-dependent movement of fluid across the ileum, hence suggesting a role as an anti-diuretic hormone (Phillips and Audsley, 1995). Subsequently, Meredith *et al*. identified the complete amino acid sequence of *Schgr*ITP, with a 72-residue mature peptide sequence and six cysteine residues proposed to participate in disulfide bridge formation (Meredith et al., 1996). The mature *Schgr*ITPL peptide is only four amino acids longer than ITP, containing a unique carboxy-terminus (Dircksen, 2009; Phillips et al., 1998). Studies have revealed that both peptides share a common N-terminal sequence, whereas the C-terminal sequences diverge significantly, thus were predicted to arise from alternative splicing (Meredith et al., 1996). Due to the shared N-terminus between the two peptide precursors, earlier studies proposed that the N-terminus permits the peptides to bind to its receptor (Phillips et al., 1998). ITP is a potent stimulator of ileal short circuit current, whereas ITP-L is devoid of such activity, suggesting an antagonistic role of ITP-L on the putative ITP receptors in the locust hindgut (Ring et al., 1998).

Differential tissue immunolocalization of ITP and ITP-L in *M. sexta* and *B. mori* revealed ITP expression in bilaterally-paired neurosecretory cells in the brain with projections to the retrocerebral complex, whereas ITP-L expression was seen in peripheral neurosecretory cells and neurons of the ventral ganglia (Dai et al., 2007). Further investigations confirmed ITP localization exclusively to the central nervous system, and ITP-L to the central nervous system and peripheral tissues (Dircksen et al., 2008; Meredith et al., 1996; Yu et al., 2016), suggesting differential functional roles for the alternatively spliced peptides. In 2007, Dai *et al*. were the first to identify a conserved ITP gene (*Itp*) in the mosquito, *Aedes aegypti*, which by alternative splicing, encodes for *Aedae*ITP-L; a longer peptide isoform with an unblocked C-terminus, and *Aedae*ITP; a shorter peptide with an amidated C-terminus. To date, the expression pattern, tissue distribution, and putative physiological function of either ITP or ITP-L has not been determined in *A. aegypti.* Herein, this study set out to characterize the tissue-specific expression and localization, as well as determine functional roles of *Aedae*ITP and *Aedae*ITP-L in the *A. aegypti* mosquito. Using a combination of molecular and physiological techniques, *Aedae*ITP and *Aedae*ITP-L was characterized in the adult stage mosquito, with expression and localization of *Aedae*ITP in the brain and the terminal ganglion while *Aedae*ITP-L was detected in the pre-terminal abdominal ganglia of the ventral nerve cord. Furthermore, using RNA interference (RNAi), the current results provide strong evidence that *Aedae*ITP and *Aedae*ITP-L play essential roles in osmotic and ionic regulation, reproductive physiology and mating behaviour in the *Aedes* mosquito. Overall, these findings advance our understanding of ITP and ITP-L neuropeptides in mosquitoes and provide novel research directions for elucidating neuropeptidergic signaling in the disease-vector, *A. aegypti*.

## Materials and Methods

### Animals

*Aedes aegypti* eggs (Liverpool strain) were collected from an established laboratory colony as described previously (Rocco et al., 2017; Sajadi et al., 2018) and hatched in double-distilled water in an incubator at 26°C on a 12:12 hour light:dark cycle. Larvae were fed a solution of 2% (w/v) brewer’s yeast and 2% (w/v) Argentine beef liver powder (NOW foods, Bloomingdale, IL, USA). For colony upkeep, female mosquitoes were fed sheep’s blood in Alsever’s solution (Cedarlane Laboratories Ltd., Burlington, ON, Canada) using an artificial feeding system (Rocco et al., 2017). Adults were provided with 10% sucrose solution *ad libidum*.

### Tissue/organ dissections, RNA extraction, and cDNA synthesis

One- and four-day old adult female (n=30) and male (n=40) *A. aegypti* were briefly anaesthetized with CO_2_ and submerged in Dulbecco’s phosphate buffered saline (DPBS; Wisent Corporation, St. Bruno, QC, Canada), and the following body segments and tissues/organs were dissected and isolated: head, thorax, midgut, Malpighian tubules (MTs), hindgut, reproductive tissues (ovaries, testes, and accessory reproductive tissues) and carcass (remaining cuticle, musculature, fat body, and abdominal ganglia). For the central nervous system expression profile, the brain, thoracic ganglia, and abdominal ganglia were collected. Whole adult RNA was obtained by collecting one- and four-day old adult female (n=10) and male (n=11) mosquitoes. For the starvation assay, whole adult male (n=6-7) and females (n=5-6) were isolated 24 h or 48 h post treatment. To confirm knockdown efficiency following double-stranded RNA (dsRNA) treatment, whole adult male (n=5-6) and female (n=5) were isolated four-, six-, and eight-days post injection. Whole adult and organ samples were stored in 1x RNA protection buffer at -20°C until further processing. Samples were then thawed at room temperature and total RNA was isolated using the Monarch Total RNA Miniprep Kit following manufacturers protocol with an on-column DNase treatment to remove genomic DNA (New England Biolabs, Whitby, ON, Canada). Purified total RNA samples were subsequently aliquoted onto a Take3 micro-volume plate and quantified on a Synergy Multi-Mode Microplate Reader (BioTek, Winooski, VT, USA). To determine *AedaeItp* and *AedaeItp-l* transcript levels, cDNA was synthesized from 500 ng (developmental expression profile), 80 ng (spatial expression profile), and 250 ng (starvation assay and dsRNA injections) total RNA using the iScript^™^ Reverse Transcription Supermix for RT-qPCR (Bio-Rad, Mississauga, ON, Canada) following manufacturers protocol, including a ten-fold dilution of cDNA following synthesis.

### RT-quantitative PCR

To measure expression profiles for *AedaeItp* and *AedaeItp-l*, transcript abundance was quantified on a StepOnePlus^™^ Real Time PCR system (Applied Biosystems, Carlsbad, CA, USA) using PowerUP^™^ SYBR® Green Master Mix (Applied Biosystems, Carlsbad, CA, USA). Cycling conditions were as follows: 1) UDG activation 50°C for 2 min, 2) 95°C for 2 min, and 3) 40 cycles of i) 95°C for 15 seconds and ii) 60°C for 1 minute. Gene-specific primers for *AedaeItp* and *AedaeItp-l* were designed over multiple exons (see **Supplementary Table S1** for list of primers) based on a previously reported mRNA sequence (Genbank Accession Numbers: (*Itp*) AY950503 and (*Itp-l*) AY950506) (Dai et al., 2007). To ensure specificity for each individual peptide-specific transcript, reverse primers for *AedaeItp* (nucleotides 418-438) and *AedaeItp-l* (nucleotides 418-428) were designed over transcript-specific exon-exon boundaries that, in the case of *AedaeItp-l*, includes exon 3 since this exon is absent in *AedaeItp* (**Supplementary Figure S1**). Relative expression levels were determined using the ΔΔC_T_ method (Livak and Schmittgen, 2001) and normalized to the geometric mean of *rp49* and *rps18* reference genes, which were previously determined as optimal endogenous controls (Paluzzi et al., 2014). Developmental expression profiles consisted of an average of 5-6 biological replicates that each included triplicate technical replicates for each target gene. Spatial expression profiles, starvation assay and dsRNA knockdown experiments consisted of 3-4 biological replicates. Primer specificity for target mRNA was assessed by conducting no reverse-transcriptase and no-template controls along with performing standard curves to calculate primer efficiencies.

### Immunohistochemistry

To examine *Aedae*ITP and *Aedae*ITP-L immunoreactivity in the central nervous system, whole one- and four-day old adult male and female *A. aegypti* were collected and incubated in freshly prepared 4% paraformaldehyde (PFA) fixative overnight at 4°C. The following day, tissues/organs dissections of the central nervous system consisting of the brain, thoracic ganglia, and the abdominal ganglia, were performed in DPBS. Tissues/organs were then permeabilized by incubating on a rocker for 1 h at RT in 4% Triton X-100 (Sigma Aldrich, Oakville, ON, Canada), 10% normal sheep serum (NSS) (v/v) and 2% BSA (w/v) prepared in DPBS, followed by three 15 min washes in DPBS. After the last wash, the DPBS was removed and substituted with a 1:1000 dilution of primary antiserum solution (0.4% Triton-X-100, 2% NSS (v/v), and 2% BSA (w/v)) in DPBS (prepared the day before use and incubated at 4°C to reduce non-specific binding) on a rocker for 96 h at 4°C. The custom primary antiserum solution was raised in rabbit against a synthetic peptide (SSFFDIECKGQFNKA) antigen corresponding to a 15-amino acid region of *Aedae*ITP and *Aedae*ITP-L (nucleotides 154-198 on common exon 2, amino acids 1-15 of the shared N-terminal sequence, **Supplementary Figure S1**), thus targeting both *Aedae*ITP and *Aedae*ITP-L (Biomatik, Kitchener, ON, Canada). Following incubation, tissues/organs were washed with DPBS four times on a rocker over the course of an hour and subsequently incubated overnight at 4°C with a goat anti-rabbit Alexa Fluor® 568 IgG (H+L) secondary antibody (Molecular Probes, Life Technologies, Eugene, OR, USA) diluted 1:200 in 10% NSS made up in DPBS and protected from light. The following day, tissues/organs were washed three times with DPBS for 15 min each. As a negative control, the anti-*Aedae*ITP/ITP-L primary antiserum was preincubated with 10 μM antigen (SSFFDIECKGQFNKA) overnight prior to use. Additionally, tissues/organs were also incubated with a no-primary control (0.4% Triton-X-100, 2% NSS (v/v), and 2% BSA (w/v) prepared in DPBS). Tissues/organs were mounted on cover slips with mounting media comprised of DPBS with 50% glycerol containing 4 μg/mL 4’6-diamidino-2-phenylindole dihydrochloride (DAPI) and were visualized on a Zeiss LSM 800 confocal laser microscope (Carl Zeiss, Jena, Germany) and processed with the Zeiss LSM Image Browser software or visualized on a Lumen Dynamics XCite^TM^ 120Q Nikon fluorescence microscope (Nikon, Mississauga, ON, Canada).

### Starvation and blood feeding assay

To examine the potential roles of *Aedae*ITP and *Aedae*ITP-L in mosquito feeding/starvation, adult males and females were isolated post-emergence and sorted into three treatment conditions: desiccated (no food or water provided), fed (10% sucrose *ad-libitum*), and starved (only water provided). The adults were collected after 24 h or 48 h, and mRNA transcript levels of *AedaeItp* and *AedaeItp-l* were examined by RT-qPCR (as described above). To investigate whether a protein-rich meal influences the transcript abundance of *AedaeItp* and *AedaeItp-l*, adult female mosquitoes (four-six day old) were given 20 min to blood feed on sheep’s blood in Alsever’s solution, and all blood fed females were subsequently isolated after 1, 6, 12 and 24 h post-bloodmeal. Blood fed females were compared to control, similarly aged (five-six day old) females that were provided sucrose *ad libitum*. Post isolation, mRNA transcript levels of *AedaeItp* and *AedaeItp-l* were examined by RT-qPCR (as described above).

### Preparation and microinjection of *AedaeItp* and *AedaeItp-l* dsRNA

Gene-specific primers were designed to amplify a region of the *AedaeItp* and *AedaeItp-l* transcripts as a template for dsRNA synthesis (ensuring no overlap with RT-qPCR primers, **Supplementary Table S1, Supplementary Figure S2**). Similar to the gene-specific probes, ds*AedaeItp* primers were designed over common exons 1 and 2, that targets and knocks down both *AedaeItp* and *AedaeItp-l* mRNA, whereas *AedaeItp-l* primers were designed over the unique exon 3 targeting only *AedaeItp-l* transcript (**Supplementary Figure S1**). The *AedaeItp* and *AedaeItp-l* target regions were amplified and cloned into pGEM-T-Easy vector and subsequently subcloned into the L4440 vector, which possesses two T7 promoters, each flanking either side of the multiple cloning site. L4440 was a gift from Andrew Fire (Addgene plasmid#1654; http://n2t.net/addgene:1654; RRID:Addgene_1654). The *AedaeItp* and *AedaeItp-l* targets were screened with a M13 forward (5’-TGTAAAACGACGCCAGT-3’) and L4440 reverse primer (5’-AGCGAGTCAGTCAGTGAGCGAG-3’) and reamplified with a T7 primer serving as a forward and reverse primer. Double stranded RNA was synthesized by *in vitro* transcription using the HiScribe T7 High Yield RNA Synthesis Kit (New England Biolabs, Whitby, ON, Canada) following manufacturers recommendations. Following synthesis, the dsRNA was incubated at 75°C for 5 min for denaturation and left at RT for 15 min to allow rehybridization, followed by RNA purification using a Monarch RNA Cleanup Kit following the manufacturer’s protocol (New England Biolabs, Whitby, ON, Canada). One-day old male and female adult mosquitoes were briefly anesthetized using CO_2_ and injected in the thorax with 1 μg of *AedaeItp*, *AedaeItp-l* or control *Egfp* (enhanced green fluorescent protein) dsRNA using a Nanoject III Programmable Nanoliter Injector (Drummond Scientific, Broomall, PA, USA).

### *In vivo* urine production assay

To determine if *Aedae*ITP and/or *Aedae*ITP-L influences urine output, adult female mosquitoes were injected with 500 nL of a HEPES buffered saline (HBS), consisting of 11.9 mM HEPES, 137 mM NaCl, and 2.7 mM KCl, titrated to a pH of 7.45 and filter sterilized before use. Based on an established protocol (Calkins and Piermarini, 2015), four-day old females were injected and placed into a graduated, packed-cell volume tube (MidSci, St. Louis, MO, USA) for two hours at 28°C with three mosquitoes per tube and excretion volumes were measured. Specifically, following the incubation period, mosquitoes were removed from the tube, which was then centrifuged at 16,000x*g* for 30 s to allow for the excreted volume to be measured visually under a dissecting microscope, via the graduated column at the bottom of the tube. Treatment females were either four-days old ds*Egfp*, ds*AedaeItp*, ds*AedaeItp-l* females (injected at one-day old), or non-dsRNA injected four-days old female mosquitoes that served as controls. Mosquito images were captured using a Zeiss Stemi 508 microscope with an Axiocam 208 color camera (Carl Zeiss, Jena, Germany).

### Mating and egg-laying assay

*A. aegypti* mosquitoes were separated at the pupal stage and individually placed into a 24-well plate to allow adults to emerge. One-day old non-mated male and female adults were then isolated and injected as follows: 1) ds*AedaeItp* or ds*AedaeItp-l* knockdown females mated with virgin males, 2) ds*AedaeItp* or ds*AedaeItp-l* knockdown males mated with virgin females, and 3) ds*AedaeItp* or ds*AedaeItp-l* knockdown females mated with ds*AedaeItp* or ds*AedaeItp-l* knockdown males. For 1), ds*AedaeItp* or ds*AedaeItp-l* females were grouped with one-day old virgin males at a 1:2 ratio of female:male per insect box (BugDorm-5 insect box, MegaView Science Co. Taiwan) between 18 and 24 h after dsRNA injection. After four-five days post-mating, knockdown females were provided a bloodmeal. For 2), ds*AedaeItp* or ds*AedaeItp-l* males were mated with two-three-day old virgin females, and females were provided a bloodmeal 48 h post-mating. Lastly, for 3), ds*AedaeItp* or ds*AedaeItp-l* males and females were mated, and females were provided a bloodmeal four-five days post injection. For all blood feeding assays, females were given 20 min to blood feed, and blood fed females were subsequently isolated and weighed individually before being placed in an inverted 25 cm^2^ cell culture flask (Corning) lined with filter paper containing 3 mL of distilled water (dH_2_O) from larvae rearing containers to promote egg laying. Laid eggs were collected after 4 days and were semi-desiccated for 72 h and counted. Females were then removed and weighed before spermathecae were dissected and viewed under a microscope to confirm insemination. Eggs were placed in 40 mL of dH_2_O with 1 mL larval food (1:1 ratio of 2% brewer’s yeast and 2% liver powder), and hatched larvae (if any) were counted after 48 h.

### Sperm Quantification

Sperm quantification in the paired seminal vesicles (male sperm storage organs) and testes of male *A. aegypti*, along with the spermathecae (sperm storage organs) of female *A. aegypti* 4 days post dsRNA injection was performed following previously published protocols (Durant and Donini, 2020; Rocco et al., 2019). The seminal vesicles and testes from male along with spermathecae from female mosquitoes (9-11 mosquitoes per dsRNA mating treatment from 3-4 mating replicates) were placed in a 96-well plate with 100 μL PBS, and gently torn open using ultrafine forceps to release spermatozoa. An additional 10 μL PBS was used to rinse the forceps and the PBS with spermatozoa was mixed thoroughly using a P100 pipette. Five 1 μL droplets of the PBS/spermatozoa mixture were spotted onto a microscope slide (previously treated with poly-L-lysine to promote sperm attachment), allowed to air dry completely, and subsequently fixed with 70% ethanol. Slides were mounted using mounting media comprised of DPBS with 50% glycerol containing 4 μg/mL DAPI, and the nuclei of spermatozoa within each 1 μL droplet was imaged under 4X magnification using an Olympus IX81 inverted microscope (Olympus Canada, Richmond Hill, ON, Canada). The nuclei of spermatozoa were counted in each 1 μL droplet, averaged across all five droplets for each animal, and multiplied by the dilution factor to determine total spermatozoa numbers within the seminal vesicle, testes, and spermatheca.

### Statistical analyses

All graphs were created and statistical analyses performed using GraphPad Prism v9.1 (GraphPad Software, San Diego, CA, USA). Data was analyzed accordingly using an unpaired t-test or one-way or two-way ANOVA with the appropriate post-hoc test as indicated in each figure caption, with differences between treatments considered significant if p<0.05.

## Results

### Tissue/organ-specific expression profile of *AedaeItp* and *AedaeItp-l* transcripts

Post-embryonic stages and selected tissues/organs of mosquitoes were examined for *AedaeItp* and *AedaeItp-l* transcript expression and compared between males and females. Developmental expression profiling revealed significant enrichment of *AedaeItp* transcript abundance in adult stage mosquitoes (**Figure 1A**), with greatest and significant enrichment in one- and four-day old adult males as well as one-day old adult females compared to fourth instar larvae. In contrast, *AedaeItp-l* transcript was significantly enriched in late pupal and adult stage mosquitoes, including one-day old males and females (**Figure 1B**). Transcript abundance of *AedaeItp* was significantly higher compared to *AedaeItp-l* abundance in adult-stage mosquitoes, with over a three-fold higher abundance of *AedaeItp* transcript compared to *AedaeItp-l* (**Figure 1C**). Additionally, *AedaeItp* transcript abundance was exclusively and significantly enriched in the head (**Figure 2A,C**) and brain (**Figure 2B,D**) in both adult male and female mosquitoes. Comparatively, expression of *AedaeItp-l* transcript was significantly enriched in the carcass (**Figure 2E,G**) and abdominal ganglia (**Figure 2F,H**) in adult mosquitoes.

**Figure 1.**
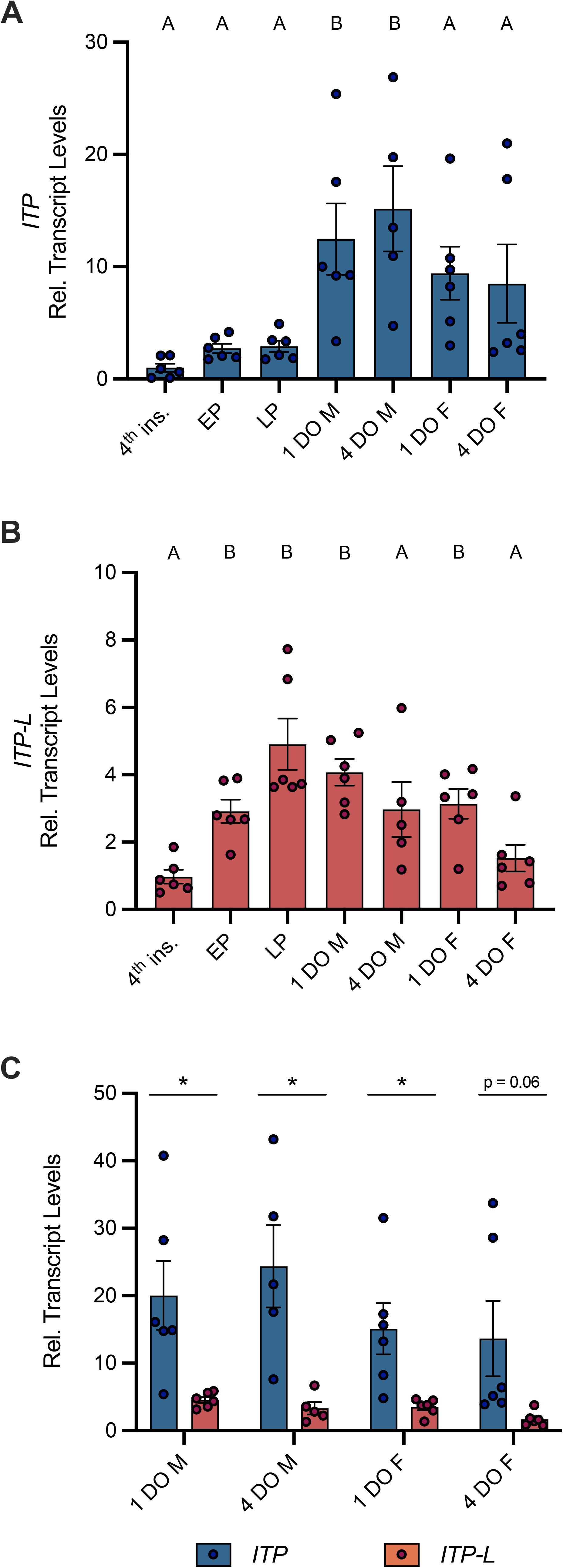
Developmental transcript expression profile of *AedaeItp* and *AedaeItp-l* in *A. aegypti.* Expression of *AedaeItp* (A) and *AedaeItp-l* (B) transcript was analyzed in post-embryonic stages of the mosquito shown relative to transcript levels in fourth instar larvae. Data labeled with different letters are significantly different from fourth instar larvae (mean±SEM; one-way ANOVA with Dunnett’s multiple comparison, p<0.05, n=5–6 biological replicates). (C) Comparison expression profile of *AedaeItp* and *AedaeItp-l* transcript levels in one and four day old adult male and female mosquitoes. Significant differences between *AedaeItp* and *AedaeItp-l* transcript levels are denoted by an asterisk (*) (mean±SEM; unpaired t-tests, p<0.05, n=5–6 biological replicates). Abbreviations: 4^th^ instar larvae (4^th^ ins), early pupa (EP), late pupa (LP), male (M), female (F), and day old (DO).

**Figure 2.**
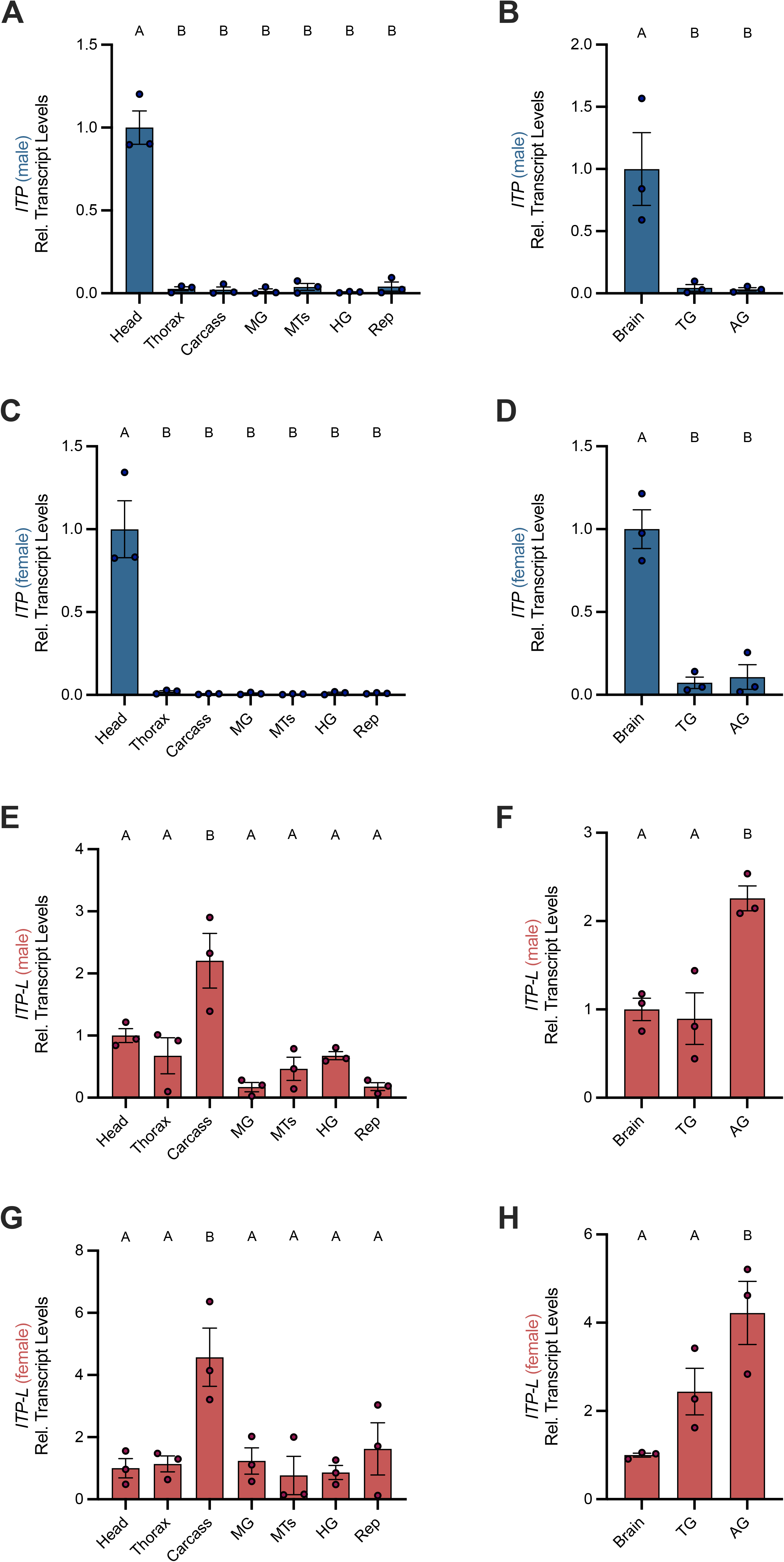
Spatial transcript expression profile of *AedaeItp* and *AedaeItp-l* in *A. aegypti.* Expression of *AedaeItp* (A–D) and *AedaeItp-l* (E–H) transcript levels were analyzed in various tissues/organs from one-day old adult males (A,B,E,F) and females (C,D,G,H) shown relative to transcript levels in the mosquito head/brain. Abbreviations: Malpighian tubules (MTs), midgut (MG), hindgut (HG), reproductive tissues/organs (Rep), thoracic ganglia (TG), and abdominal ganglia (AG). Bars labeled with different letters are significantly different from head/brain (mean±SEM; one-way ANOVA with Dunnett’s multiple comparison, p<0.05, n=3 biological replicates).

### *Aedae*ITP- and *Aedae*ITP-L-like immunoreactivity in the central nervous system

Using whole mount immunohistochemistry, the central nervous system from one-day old male and female mosquitoes revealed *Aedae*ITP- and *Aedae*ITP-L-like immunostaining in three pairs of lateral neurosecretory cells in the medial anterior region of each brain hemisphere with axonal processes projecting anteriorly near an additional single pair of lateral neurosecretory cells (**Figure 3A-E**). A number of varicosities and blebs can be seen peripherally near the axonal projections, suggestive of release sites of these neuropeptides (**Figure 3A-E**). In abdominal ganglia 2-6, *Aedae*ITP- and *Aedae*ITP-L-like staining was observed in a single pair of neurosecretory cells, positioned laterally on either side of each ganglia (**Figure 3F,G**). The terminal ganglion, which is a fusion of abdominal ganglia 7 and 8, revealed *Aedae*ITP- and *Aedae*ITP-L-like immunostaining in a single pair of neurosecretory cells located in the anterior region of the ganglion (corresponding to abdominal ganglia 7) (**Figure 3H**). Staining in these cells and projections were absent in control treatments where either the antiserum was preabsorbed with the ITP antigen or the omission of the primary antiserum (**Supplementary Figure S1**).

**Figure 3.**
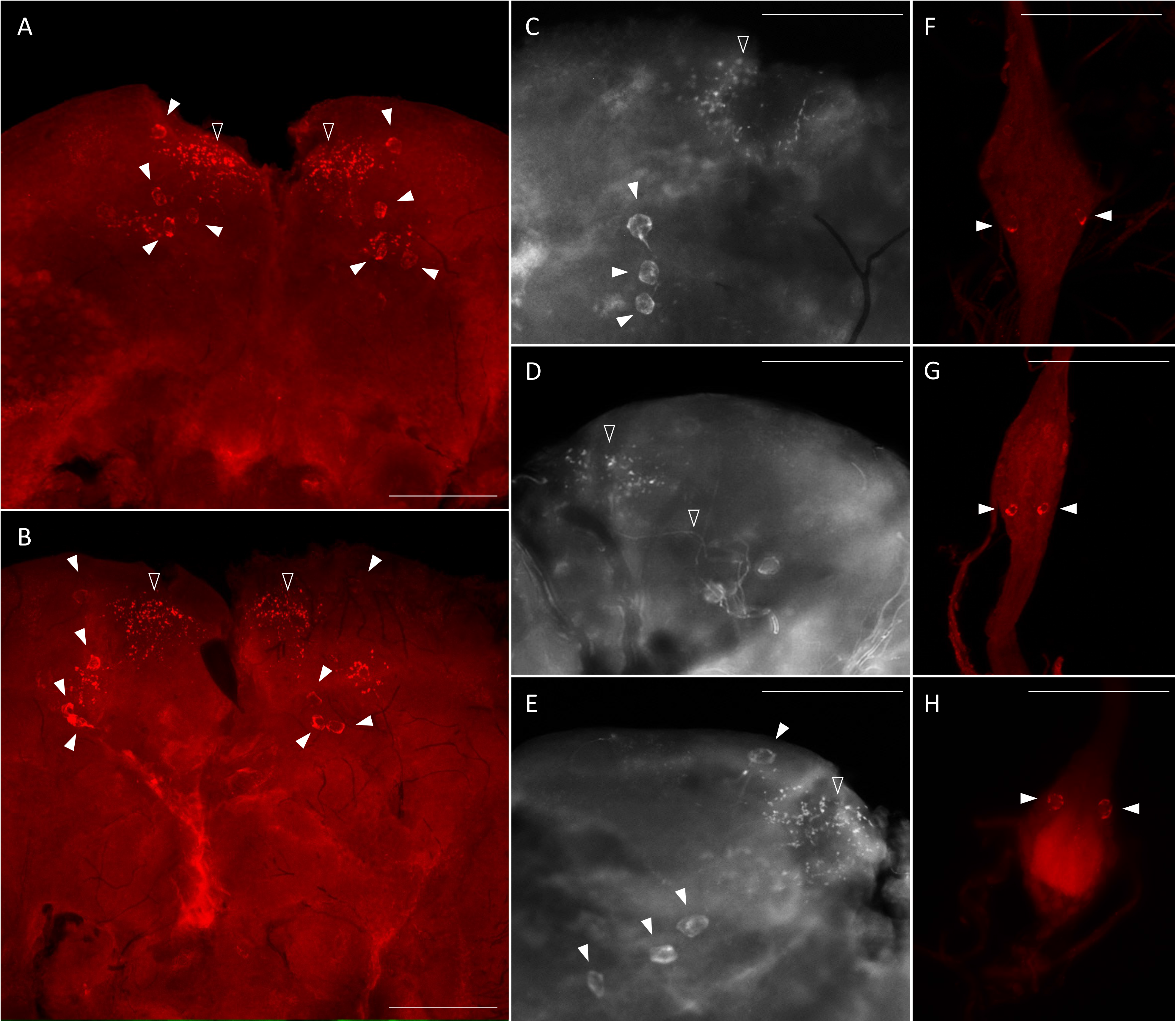
Immunolocalization of *Aedae*ITP and *Aedae*ITP-L in the central nervous system of the *A. aegypti* mosquito. *Aedae*ITP- and *Aedae*ITP-L-like immunoreactivity was examined in (A) male and (B) female brains, in four pairs of neurosecretory cells (indicated by white arrowheads), with axonal processes projecting anteriorly (C-E), towards varicosities and blebs on the periphery of the brain (indicated by empty arrowheads). (F,G) Ventral view of abdominal ganglia showing a single pair of lateral neurosecretory cells and (H) an anterior pair observed in the terminal ganglion. Scale bars: (A-B) 200 μM; (C-H) 100 μM.

### Roles of *Aedae*ITP and *Aedae*ITP-L in desiccation and starvation stress and blood feeding

Post emergence, one-day old adult male and female mosquitoes were placed in either a fed (provided a 10% sucrose meal), desiccated (no food or water provided), or starved (provided a water meal, no sucrose) condition and isolated after 24 h or 48 h to examine a potential role of *Aedae*ITP and *Aedae*ITP-L in desiccation and starvation stress (**Figure 4A**). When adult male and female mosquitoes were placed in the starved or desiccated treatment for 24 h, there was no difference in mRNA abundance of either *AedaeItp* or *AedaeItp-l* compared to control fed conditions (**Figure 4B,D,F,H**). However, when the mosquitoes were subjected to these treatments for 48 h, there was a significant enrichment of both *AedaeItp* and *AedaeItp-l* transcript levels in the desiccated condition (ranging between ∼1.75 and ∼3.5-fold) compared to control fed animals (**Figure 4C,E,G,I**) while no change to the transcript levels in animals that were starved but provided with water. Next, to determine if *Aedae*ITP and/or *Aedae*ITP-L may play a role in relation to blood feeding, four-to six-day old adult female mosquitoes were provided a bloodmeal to examine whether a protein-rich meal influences the transcript abundance of *AedaeItp* and *AedaeItp-l* (**Figure 5A**). *AedaeItp* mRNA abundance did not change significantly compared to control, sucrose-fed females over any of the measured timepoints between 1 and 24 h post-blood feeding (**Figure 5B**) although abundance trended lower at the 1, 6 and 12 hr post-blood feeding timepoints. Similarly, *AedaeItp-l* mRNA abundance did not change significantly at any of the post blood-feeding timepoints in comparison to control, sucrose-fed females (**Figure 5C**).

**Figure 4.**
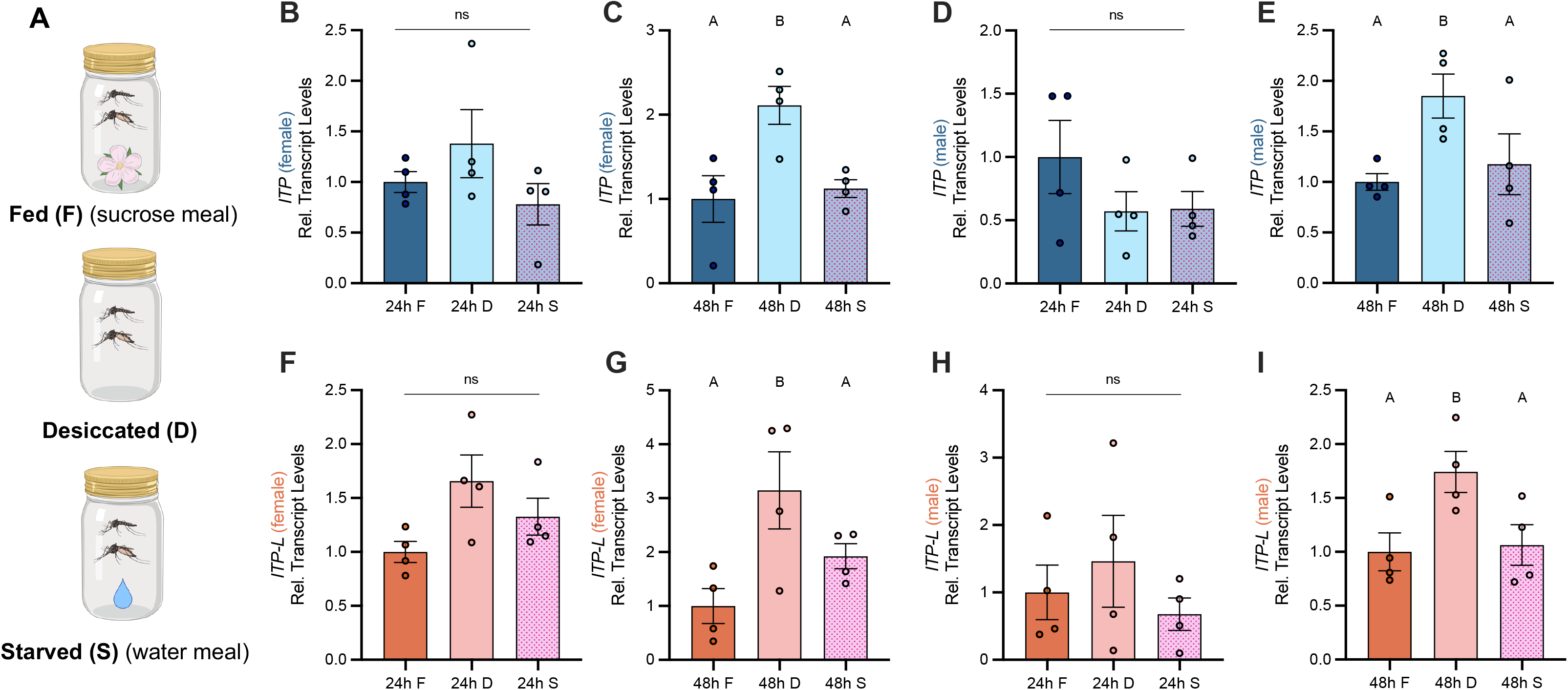
Effect of desiccation and starvation stress on transcript levels of *AedaeItp* and *AedaeItp-l* in adult *A. aegypti*. (A) Post-emergence, adult males were placed in a fed (sucrose meal provided), desiccated (no food or water provided), and starved (only water provided) condition for 24 h and 48 h, and abundance of (B-E) *AedaeItp* and (F-I) *AedaeItp-l* transcript were analyzed, shown relative to transcript levels in the control (fed) adults. Abbreviations: fed (F), desiccated (D), and starved (S). Bars labeled with different letters are significantly different from the 24 h fed adult controls (mean±SEM; one-way ANOVA with Bonferroni multiple comparison, p<0.05, n=4 biological replicates).

**Figure 5.**
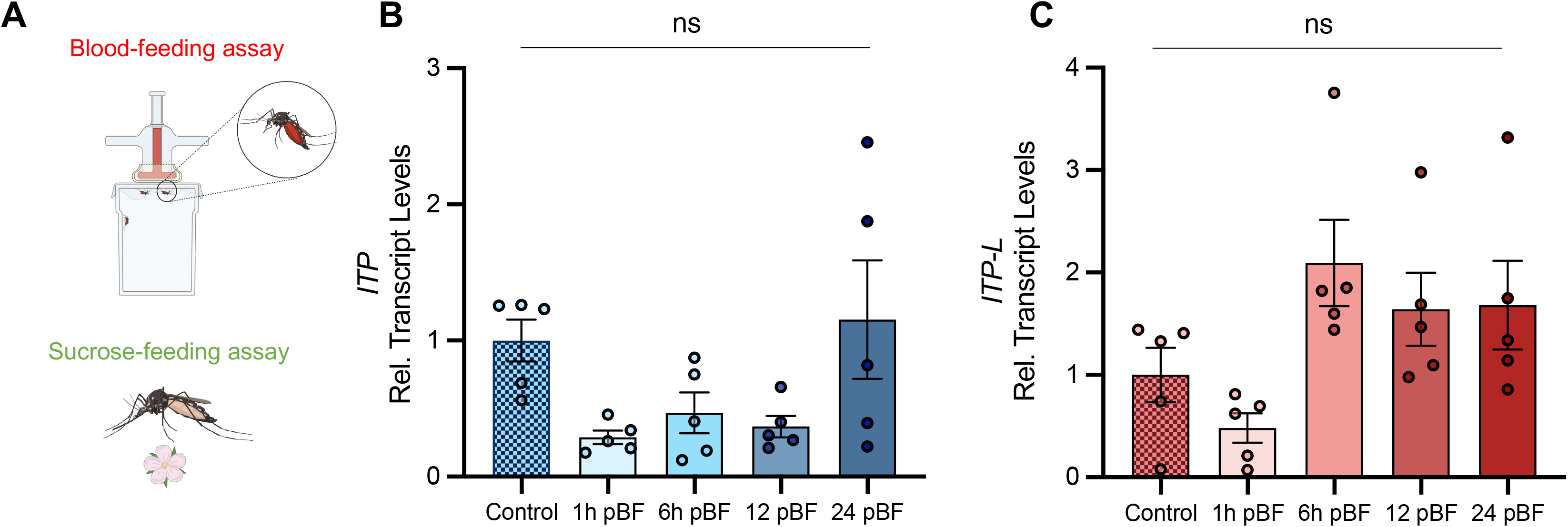
Effect of blood feeding on transcript levels of *AedaeItp* and *AedaeItp-l* in adult *A. aegypti*. (A) Four-to six-day old adult females were blood fed and isolated at 1, 6, 12, and 24 post-blood feeding and (B) *AedaeItp* and (C) *AedaeItp-l* transcript levels were analyzed, shown relative to control, sucrose-fed females. Abbreviations: pBF (post-blood feeding). No significance (ns) reflects comparisons with control, sucrose-fed females (mean±SEM; one-way ANOVA with Tukey’s multiple comparison, p<0.05, n=5 biological replicates).

### dsRNA knockdown of *AedaeItp* and *AedaeItp-l* in adult *Aedes* mosquitoes

RNA interference of *AedaeItp* and *AedaeItp-l* expression was accomplished through dsRNA-injections of one-day old adult male and female mosquitoes (**Supplementary Figure S3A**). Given the conserved exons 1, 2, and 4 between *AedaeItp* and *AedaeItp-l*, the ds*AedaeItp* primers were designed over a common exon 2 resulting in the knockdown of both *AedaeItp* and *AedaeItp-l* transcripts. However, ds*AedaeItp-l* primers were designed over the unique exon 3 allowing knockdown of only the *AedaeItp-l* transcript. Relative to ds*Egfp*-injected control mosquitoes, *AedaeItp* and *AedaeItp-l* transcripts were significantly reduced by ∼75% in four-day old male (**Supplementary Figure S3B**) and female (**Supplementary Figure S3C**) mosquitoes injected with ds*AedaeItp.* Comparatively, ds*AedaeItp-l* treatment resulted in a significant decrease in *AedaeItp-l* transcript abundance by ∼80% in males (**Supplementary Figure S3B**) and ∼60% in females (**Supplementary Figure S3C**) in four-day old adults, whereas *AedaeItp* transcript abundance was unaffected. To confirm injection alone does not influence *AedaeItp* and *AedaeItp-l* transcript levels, four-day post-ds*Egfp* injected animals were compared to four-day old non-injected mosquitoes (**Supplementary Figure S3D**), with no significant changes in *AedaeItp* and *AedaeItp-l* transcript abundance in ds*Egfp*-injected males and females compared to non-injected mosquitoes. *AedaeItp* and *AedaeItp-l* transcript restored to normal levels within six and eight days post-injection in males and eight days post-injection in females (**Supplementary Figure S3E-H**).

### dsRNA knockdown confirms *Aedae*ITP and *Aedae*ITP-L immunolocalization

To further confirm ds*AedaeItp*- and *AedaeItp-l* knockdown and differentiate between *Aedae*ITP and *Aedae*ITP-L immunolocalization, staining in the CNS was examined in mosquitoes four-days post-dsRNA injection. Wholemounts of ds*AedaeItp*-injected male and female mosquitoes showed no *Aedae*ITP- and *Aedae*ITP-L-like immunostaining in the brain (**Supplementary Figure S4A,B**), abdominal ganglia (**Supplementary Figure S4C**), and in the terminal ganglion (**Supplementary Figure S4D**). In contrast, ds*AedaeItp-l*-injected mosquitoes did not exhibit changes to immunostaining in the brain (**Supplementary Figure S4E,F**), with similar staining as described above (**Figure 3A-E**), and in control ds*Egfp*-injected mosquitoes (**Supplementary Figure S5A,B**). However, as expected, knockdown of *AedaeItp-l* resulted in abolished staining of neurosecretory cells in the pre-terminal abdominal ganglia (**Supplementary Figure S4G**), and interestingly, no change to the immunostaining in the terminal ganglion (**Supplementary Figure S4H**), with a single pair of immunoreactive cells as described above (**Figure 3H**) and also observed in control ds*Egfp*-injected mosquitoes (**Supplementary Figure S5C,D**).

### *AedaeItp* knockdown influences urine output in adult females

To determine if *Aedae*ITP and/or *Aedae*ITP-L influences ionoregulation and urine excretion, we volume loaded ds*AedaeItp* and ds*AedaeItp-l* injected females with saline and measured their urine output over two hours post volume loading. Four-day old HBS-injected females excreted a volume of 429.5±53.49 nL of urine, which was significantly higher (∼4-fold) compared to control four-day old non-HBS injected females (106.8±15.39 nL) (**Figure 6A**). Notably, ds*Egfp* treated females excreted a similar volume of urine (391.6±61.03 nL) compared to HBS-injected females. In contrast, ds*AedaeItp* mosquitoes injected with HBS secreted a significantly higher amount of urine, 972.7±70.18 nL, approximately 2.5-fold higher compared to ds*Egfp*-injected females. Interestingly, no significant change in urine output was observed in ds*AedaeItp-l* females injected with HBS (604.2±78.87 nL), compared to HBS loaded control (non-dsRNA injected) and ds*Egfp*-injected females. The overall effect of ds*AedaeItp* and ds*AedaeItp-l* injections on urine output was studied by examining abdomen distension immediately after and two hours post-HBS injection. Two hours post-volume loading, a less distended abdomen was observed in ds*AedaeItp*-injected females (**Figure 6H– I**) compared to control ds*Egfp*- and HBS-injected females (**Figure 6B–G**). Comparatively, a moderately distended abdomen was observed in ds*AedaeItp-l*-injected females (**Figure 6J–K**) two hours post saline-injection.

**Figure 6.**
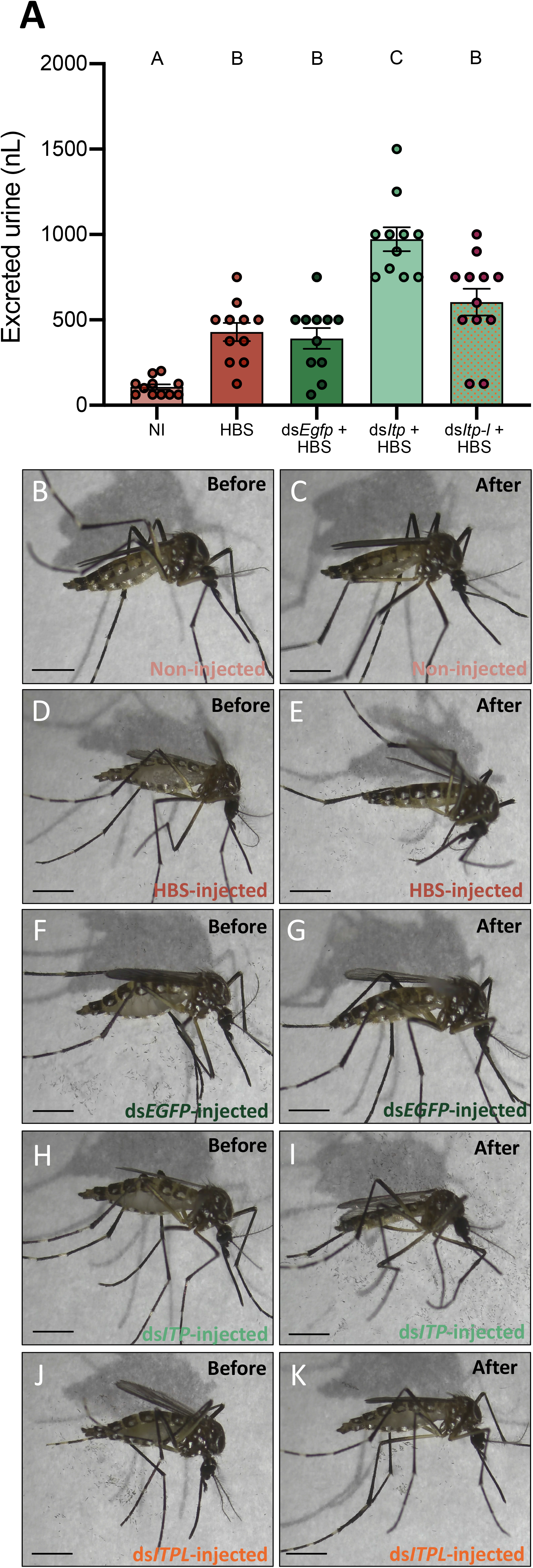
Effect of *AedaeItp* and *AedaeItp-l* knockdown on urine excretion in adult female *A. aegypti*. (A) Four-day old females were injected with 500 nL of a HEPES buffered saline (HBS) and allowed to excrete for two hours. Bars labeled with different letters are significantly different from each other (mean±SEM; one-way ANOVA with Bonferroni multiple comparison, p<0.05, n=11–12). (B-K) Images of female mosquitoes immediately following and two hours post-HBS injection. Scale bars: 1 mm.

### *AedaeItp* and *AedaeItp-l* knockdown influences male and female reproductive success

To assess the roles of *Aedae*ITP and *Aedae*ITP-L in reproductive behaviour and physiology, dsRNA-injected virgin mosquitoes were placed into one of the following mating combinations: 1a) ds*AedaeItp* or 1b) ds*AedaeItp-l* females mated with control, non-dsRNA males, 2a) ds*AedaeItp* or 2b) ds*AedaeItp-l* males mated with control, non-dsRNA females, and 3a) ds*AedaeItp* or 3b) ds*AedaeItp-l* females mated with ds*AedaeItp* and ds*AedaeItp-l* males. When ds*AedaeItp* females were mated with normal males, there was a significant reduction in the incidence of blood feeding (**Figure 7A**), reduced bloodmeal engorged by the female (**Figure 7B**), reduction in the number of eggs laid (**Figure 7C**), and an overall reduction in the percentage of larvae hatching per female (**Figure 7D**) was observed. More specifically, there was a ∼90% reduction in preference for blood feeding by ds*AedaeItp* females and ∼75% reduction in ds*AedaeItp-l* females (**Figure 7A**) with both treatments leading to a reduction in the bloodmeal volume imbibed (**Figure 7B**). Females treated with ds*AedaeItp-l* oviposited a similar number of eggs and had a comparable percentage of hatched larvae as control ds*Egfp*-females while ds*AedaeItp* caused a drastic reduction (∼50%) in eggs oviposited by females and a more dramatic impact on larval hatching success (**Figure 7C,D**). In males injected with either ds*AedaeItp* or ds*AedaeItp-l* and mated with normal females, this did not influence female preference for blood feeding or the volume of blood imbibed (**Figure 7E-F**). However, normal females mated with ds*AedaeItp* and ds*AedaeItp-l* injected males had significantly reduced number of eggs oviposited (**Figure 7G**), as well as a reduced percentage of viable eggs and larvae hatching per female (**Figure 7H**). Interestingly, ds*AedaeItp* and ds*AedaeItp-l* females mated with ds*AedaeItp* and ds*AedaeItp-l* males almost completely abolished the preference for blood feeding (**Figure 7I**) with the few that did blood feed imbibing a significantly lower blood volume (**Figure 7J**), with almost no eggs laid by the female (**Figure 7K**), and complete absence of larval hatching (**Figure 7L**). Notably, the reduced preference for blood feeding limited the number of blood fed females used for subsequent studies. No major changes were observed in the weight of blood fed females post-egg laying (4 days post-feeding) in any of the treatment regimens (**Figure 7B,F,J**), indicating long term volume balance is not impacted by knockdown of *AedaeItp* or *AedaeItp-l* although there was a small (but significant) increase in the weight of *AedaeItp-l* injected females (**Figure 7B**). Mated males and females injected with control ds*Egfp* had similar preference for blood feeding, weight of blood fed females, produced similar number of eggs and comparable larval hatching as non-injected females (**Supplementary Figure S6**).

**Figure 7.**
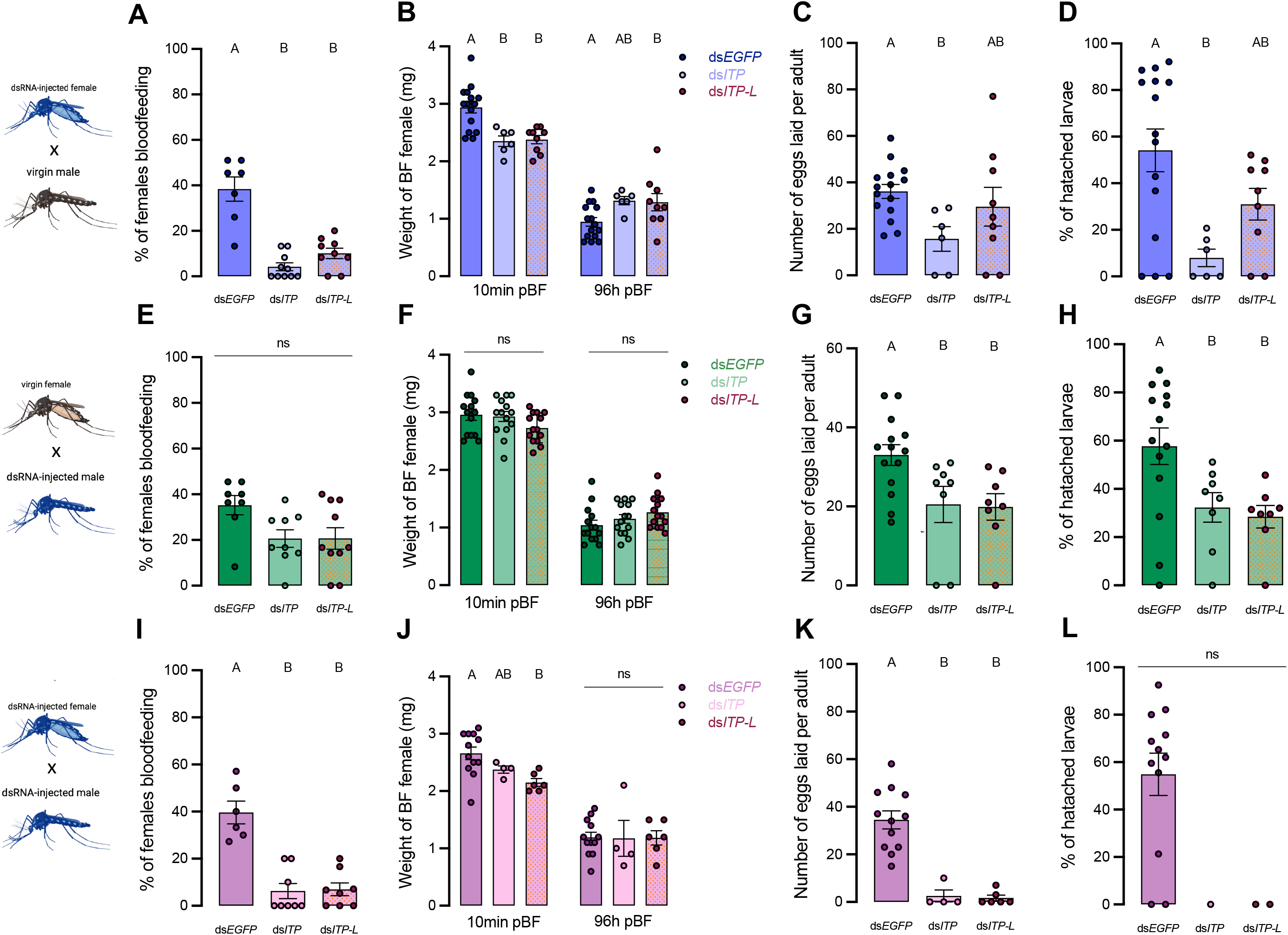
Effect of *AedaeItp* and *AedaeItp-l* knockdown on blood-feeding, egg laying, and larval hatching (egg viability) in adult *A. aegypti*. (A-D) ds*AedaeItp* or ds*AedaeItp-l* females mated with normal males, (E-H) ds*AedaeItp* or ds*AedaeItp-l* males mated with normal females, and (I-L) ds*AedaeItp* or ds*AedaeItp-l* females mated with ds*AedaeItp* or ds*AedaeItp-l* males. The effect of ds*AedaeItp* or ds*AedaeItp-l* knockdown was tested on (A,E,I) preference for blood feeding, (B,F,J) weight of blood fed female before and after egg collection, (C,G,K) number of eggs oviposited, and (D,H,L) percentage of larval hatching. Bars labeled with different letters are significantly different from control, ds*Egfp* injected age/time-matched adults (mean±SEM; one-way ANOVA with Bonferroni multiple comparison, p<0.05, (A,E,I, n=6–11 mating replicates, each point represents individual replicate values) (B-D,F-H,J,K, n=1–15, each point represents data from individual females), (ns denotes no statistical significance).

### *AedaeItp* and *AedaeItp-l* knockdown reduces spermatozoa count

In light of the results above, it supports the notion that knockdown of *AedaeItp* and *AedaeItp-l* may have a distinct role in male reproductive biology separate from their effects on females since pairings between normal females and knockdown males revealed females had normal preference for blood feeding as well as bloodmeal weight; however, oviposition rates by females and larval hatching rates were significantly reduced. Considering the overall reduced preference for blood feeding in ds*AedaeItp* and ds*AedaeItp-l* females, we speculated that *Aedae*ITP and *Aedae*ITP-L may play an essential role in spermatozoa production and release. Consequently, sperm was collected separately from the testes and seminal vesicles of four-day old ds*AedaeItp*- and ds*AedaeItp-l*-injected males (from all three mating conditions described above) and spermathecae of four-day old ds*AedaeItp*- and *AedaeItp-l*-injected females (again, from all three mating conditions noted above), and the quantity of mature spermatozoa was compared to ds*Egfp*-injected animals (**Figure 8A-B**). For spermatozoa collected from the seminal vesicles, ds*AedaeItp*- and ds*AedaeItp-l*-injected females mated with normal males produced similar mature spermatozoa counts compared to control ds*Egfp* animals (**Figure 8C, Supplementary Figure S7A-B,E**). Interestingly, knockdown resulted in significant reductions in the number of spermatozoa in the seminal vesicle, from both ds*AedaeItp*- and ds*AedaeItp-l*-injected males mated with control females and when mated with ds*AedaeItp*- and ds*AedaeItp-l* injected females (**Figure 8C, Supplementary Figure S7C,D,F,G**). Similar trends were observed for the number of spermatozoa collected directly from the male testes (**Figure 8C, Supplementary Figure S7H-N**). Comparatively, *AedaeItp*- and *AedaeItp-l* knockdown resulted in an ∼85% reduction in mature spermatozoa counts in the spermatheca in all three mating conditions (**Figure 8C, Supplementary Figure S7O-U**). Animals injected with ds*Egfp* resulted in similar number of spermatozoa in the testes, seminal vesicles, and spermathecae compared to non-injected animals (**Supplementary Figure S8**).

**Figure 8.**
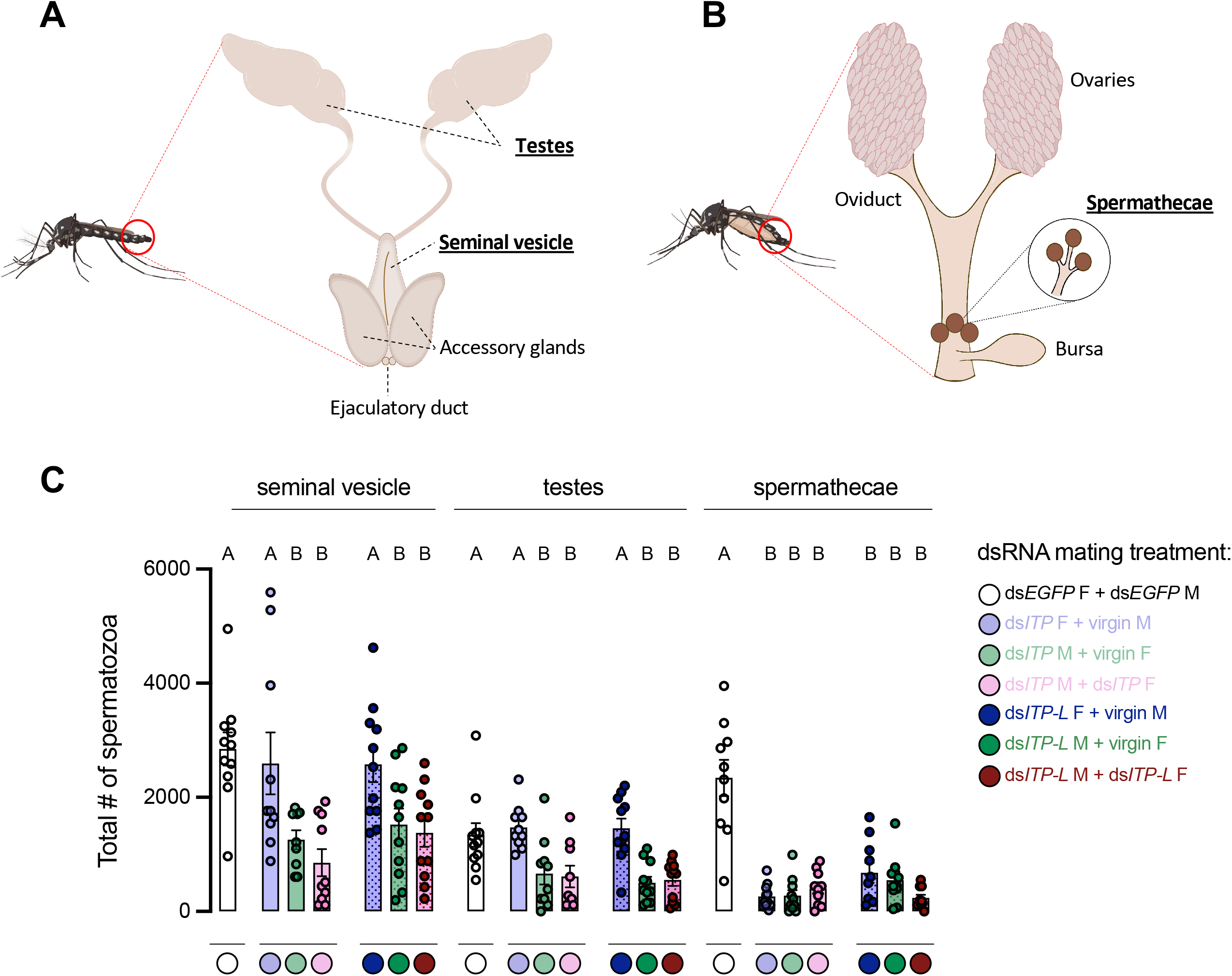
Total number of spermatozoa in the male testes and seminal vesicle along with female spermathecae of adult *A. aegypti* following RNAi (dsRNA)-mediated knockdown of *AedaeItp* or *AedaeItp-l*. Schematic diagram showing the (A) male and (B) female reproductive system of the adult mosquito. (C) Total spermatozoa number within the male testes and seminal vesicle, and female spermathecae of adults four-days after ds*AedaeItp* or ds*AedaeItp-l* injection. Abbreviations: male (M) and female (F). Bars labeled with different letters are significantly different from the organ-specific (seminal vesicle, testes and spermathecae) number of spermatozoa in control, ds*Egfp* injected adults (mean±SEM; one-way ANOVA with Bonferroni multiple comparison, p<0.05, n=9–11, each point represents individual replicate values).

## Discussion

ITP and ITP-L belong to the CHH family of neuropeptides and have been functionally characterized in many insect species (Begum et al., 2009; Dircksen et al., 2008; Drexler et al., 2007; Gáliková et al., 2018; Nagai et al., 2014; Sun et al., 2020; Webster et al., 2012; Yu et al., 2016). However, while several studies have examined ITP signaling pathways in insects, ITP/ITP-L receptors have generally not been identified and characterized thus far, except for in the domestic silk moth *B. mori* (Nagai et al., 2014). In a recent study using *Drosophila melanogaster* as a conserved tumour model, isoform F of ion transport peptide (ITPF) was found to be secreted by gut tumour cells and acts as an antidiuretic hormone targeting the tachykinin receptor (TkR99D) in Malpighian tubules leading to compromised renal function that results in the accumulation of excess fluid (Xu et al., 2023). This latest report is an intriguing considering tachykinins have been reported as diuretic factors in a number of insects (see Nässel et al., 2019) including recently in *D. melanogaster* (Agard *et al*., 2024).

In the present study, the *A. aegypti* ITP and ITP-L peptides have been localized, confirming the distribution of these peptides in neurosecretory cells and processes within the mosquito central nervous system. Additionally, prospective physiological functions have been investigated for *Aedae*ITP and *Aedae*ITP-L including roles in feeding, urine output, and reproductive success of adult male and female mosquitoes. This is the first report that examines the distribution, localization, and physiological function of the ITP/ITP-L signaling system in *A. aegypti* mosquitoes.

### Distribution pattern of *Aedae*ITP and *Aedae*ITP-L in the CNS

Expression profiles of transcripts encoding *A. aegypti* ITP and ITP-L were measured to reveal potential functional or sex-specific roles for these peptides. Examination of the developmental and tissue-specific expression profile revealed enrichment of *AedaeItp* and *AedaeItp-l* in both one- and four-day old male and females, with significant enrichment of *AedaeItp* in the brain and *AedaeItp-l* in the abdominal ganglia. The expression and distribution of ITP and ITP-L in the central and peripheral nervous system has been examined in numerous insects (Dai et al., 2007; Dircksen et al., 2008; Meredith et al., 1996; Yu et al., 2016). While *Itp* mRNA expression has been detected only in the nervous system, evidence has suggested *Itp-l* expression in peripheral tissues as well. In *T. castaneum*, *Itp-l* transcript expression was found to be highly expressed in the midgut (Begum et al., 2009), and in the Malpighian tubules and hindgut in *S. gregaria* (Meredith et al., 1996).

Previous studies in *M. sexta* revealed that *Mas*ITP and *Mas*ITPL are differentially expressed in mainly nonoverlapping populations of central and peripheral neurons, which includes neuronal projections from the CNS (Dai et al., 2007). RT-PCR, immunohistochemistry, and *in situ* hybridization studies indicated expression of *Mas*ITP exclusively in the brain where it was localized to two neuron types; in type Ia_2_ neurosecretory cells, with axonal projections to the retrocerebral complex (Copenhaver and Truman, 1986; Homberg et al., 1991; Zitnan et al., 1995; Zitnan and Adams, 2005) and in small neurons adjacent to type Ia_2_ cells, established as interneurons since their projections remain within the protocerebrum (Dai et al., 2007). Thus, in *M. sexta*, it was suggested that ITP is released as a neurohormone from type Ia_2_ cells into the haemolymph, whereas ITP produced in the small interneurons may serve transmitter or modulatory functions in the brain (Dai et al., 2007). Similarly in *S. gregaria*, ITP is believed to be synthesized in neurosecretory cells of the pars intercerebralis of the brain where it is then transported for storage and eventual release in the corpora cardiaca (Meredith et al., 1996). The existence of ITP-L transcripts was first reported in *S. gregaria* (Macins et al., 1999), while the mature peptide was identified by Dai *et al*., (2007) demonstrating ITP-L transcript and peptide distribution in the central and peripheral nervous system of insects, including *B. mori*, *M. sexta*, and the grasshopper, *Schistocerca americana*. Relatively weak ITP-L immunoreactivity was observed in the brain type Ia_2_ cells but was found to be completely absent in axons and terminals within the retrocerebral complex (Dai et al., 2007). ITP-L peptides are abundant in the ventral ganglia, flight muscle, MTs, and ileal tissues, indicating a possible distinctive function from ITP (Meredith et al., 1996; Phillips and Audsley, 1995).

Through RNAi-mediated knockdown, we confirmed *Aedae*ITP immunoreactivity in at least four pairs of neurosecretory cells in the anterior region of the protocerebrum and in a single pair of lateral neurosecretory cells in the terminal ganglion. In contrast, *Aedae*ITP-L immunoreactivity was observed in one pair of lateral neurosecretory cells in each abdominal ganglia of the ventral nerve cord. In general, expression patterns of *AedaeItp* and *AedaeItp-l* were similar to those described in *M. sexta*, *T. castaneum*, *B. mori*, and *D. melanogaster* (Begum et al., 2009; Dai et al., 2007; Dircksen, 2009; Dircksen et al., 2008). In *T. castaneum*, *Itp* expression was in five pairs of brain cells on the dorsal side of the protocerebral hemispheres and in a pair of cells in the abdominal terminal ganglion (Begum et al., 2009). Similarly, ITP expression was observed in four pairs of brain cells in *D. melanogaster*. While still inconclusive, previous immunohistochemical studies in other insects allude to protocerebral cells with projections to the corpora cardiaca and allata, and the cells in the terminal abdominal ganglion may have projections to the hindgut (Dircksen et al., 2008) and possibly to reproductive organs (Begum et al., 2009), which provide insight on novel functions for the ITP/ITP-L signaling system. Given the immunoreactivity of *Aedae*ITP in the terminal ganglion, this suggests potential iono-regulatory roles in the hindgut, acting possibly as an anti-diuretic hormone to increase water or ion reabsorption, similar to activity seen in the desert locust (Audsley et al., 1992a; Audsley et al., 1992b).

### Roles of ITP and ITP-L in feeding and urine excretion

The challenges of osmotic and ionic regulation vary between distinct environmental conditions that the adult mosquito might encounter. In desiccating environments, insects must safeguard water balance and reduce the rate of water loss (Terhzaz et al., 2012). The current findings reveal an increase in both *AedaeItp* and *AedaeItp-l* transcript levels after 48 hours of combined desiccation and starvation stress in both adult male and female mosquitoes, which was not observed in animals undergoing starvation stress alone. These findings corroborate data reported in *Drosophila*, where ITP was linked as a natural component of desiccation and osmotic stress responses, since both stressors triggered an increase in *Itp* expression, while ITP knockdown reduced survival under desiccation and osmotic stress (Gáliková et al., 2018). *Droso*ITP plays roles in hunger, thirst, and excretion in *Drosophila* suggesting that ITP-regulated changes to physiology and behaviour represent critical insect responses to cope with reduction in body water (Gáliková et al., 2018).

The discovery of the first anti-diuretic hormone mediating its effects on the insect hindgut was described by Audsley *et al*., (1992a) when ITP was purified from the corpora cardiaca of the locust, *S. gregaria*. A conserved *Itp* gene was later uncovered in the genome of the mosquito, *A. aegypti,* raising the prospect for a similar role in maintaining iono- and osmo-regulation (Dai et al., 2007). *A. aegypti* mosquitoes are reliant on an efficient excretory system comprised of the MTs and hindgut (Coast, 2007), functioning to counter disturbances to their haemolymph. The MTs, which are functional analogs of vertebrate kidneys, are responsible for the formation of primary urine (Coast, 2007), driven by the V-type H^+^-ATPase (Wieczorek, 1992) that permits transport of Na^+^ and K^+^ cations across the membrane (Beyenbach, 2003) via a putative H^+^/cation exchanger (Wieczorek et al., 2009). The MTs are regulated by various diuretic and anti-diuretic hormones, which in *A. aegypti* includes the biogenic amine 5-hydroxytryptamine (5HT) (Clark and Bradley, 1998; Veenstra, 1988), DH_31_ (Coast et al., 2005), DH_44_ (Clark et al., 1998; Clark et al., 1998), kinin-like peptides (Lu et al., 2011; Pietrantonio et al., 2005) and CAPA that inhibits the activity of select diuretic hormones (Sajadi et al., 2018; Sajadi et al., 2020; Sajadi et al., 2023). The primary urine then enters the hindgut, where it is further modified through secretory and reabsorptive processes. Here, we show that *Itp* knockdown (but not *Itp-l* knockdown) leads to increased excretion of urine, supporting a possible anti-diuretic role for *Aedae*ITP. Thus, it suggests that *Aedae*ITP promotes water reabsorption in the hindgut similar to mechanisms described in *S. gregaria* where *Schgr*ITP was found to stimulate chloride-dependent water reabsorption in the ileum, promoting an increase in Na^+^, K^+^ and Cl^-^ transport (Audsley et al., 1992b; Audsley et al., 1992a) Interestingly, while *Schgr*ITP promotes a reabsorptive role on ileal tissue, ITP-L did not display any stimulatory effect, instead inhibiting the stimulatory effect of synthetic ITP (Ring et al., 1998). A key step to understanding the ITP and ITP-L actions in the *Aedes* mosquito is to identify the as yet unknown *A. aegypti* ITP receptor. The first presumed receptors for ITP and ITPL were characterized in the silkworm *Bombyx mori* (Nagai et al., 2014). Specifically, Nagai *et al*., (2014) identified three *B. mori* orphan GPCRs as receptors for ITP and ITP-L, that all responded to recombinant ITP, with elevating levels of intracellular cGMP upon receptor binding (Nagai et al., 2014), which support the suggested ITP’s mode of action on ileal ion transport involving this second messenger in *S. gregaria* (Audsley et al., 2013). In the locust, *Schgr*ITP is proposed to bind to two different receptors, a G-protein coupled receptor and a membrane bound guanylate cyclase, on the ileal basolateral membrane, increasing both cyclic GMP (cGMP) and cyclic AMP (cAMP) levels, to regulate ion and fluid transport (Audsley et al., 2013). cGMP stimulates Cl^-^ reabsorption and H^+^ secretion across the ileum, whereas cAMP stimulates Na^+^, K^+^, and Cl^-^ reabsorption (Audsley et al., 2013). To further understand how these second messengers facilitate the physiological actions of *Schgr*ITP on the ileum will require the endogenous ITP receptor(s) to be characterized.

### Role of ITP and ITP-L in reproductive behaviour and success

Female *A. aegypti* are day-biting mosquitoes, taking a single or multiple bloodmeals to obtain vitamins, nutrients, proteins, and minerals for egg development (Beyenbach, 2003). Transcript levels of *AedaeItp-l* and *AedaeItp* levels remained unchanged over the time points we examined post blood feeding. Nonetheless, to elucidate whether ITP and ITP-L signaling might be involved in mosquito reproductive biology, RNAi was utilized to knockdown expression of *Itp* and *Itp-l* in adult *A. aegypti.* Overall, *AedaeItp* and *AedaeItp-l* knockdown female mosquitoes had a lower preference for blood feeding, laid fewer eggs, and had significantly reduced larval hatching. In *T. castaneum*, ITP and ITP-L are required throughout all life stages and are essential for reproduction and offspring survival (Begum et al., 2009). Knockdown of both ITP and ITP-L resulted in dramatic decreases in egg numbers and in survival of eggs, with reduced ovaries that lack mature ovarioles in the ITPL knockdown females (Begum et al., 2009). In contrast, ITP knockdown females had fully developed ovaries, however showed reduced oviposition rates and offspring survival. These developmental defects in *T. castaneum* were suggested to be due to hormonal imbalance in ovarian development, or indirectly caused by mating deficiencies, preventing exposure to male ejaculatory products essential for completion of ovarian development (Begum et al., 2009).

Considering the finding that *AedaeItp* and *Itp-l* knockdown mosquitoes resulted in fewer eggs laid by females mated with knockdown males, we predicted that transfer of sperm or sperm storage may be targeted. The regulation and entry of sperm into, protection within, and release from the storage organs (seminal vesicle in males, and spermathecae in females) requires both male and female-derived molecules (Avila et al., 2010; Avila et al., 2015; Schnakenberg et al., 2011). In male mosquitoes, spermatogenesis occurs in the paired testes, allowing for mature sperm cells, spermatozoa, to be synthesized (Rocco et al., 2017; Rocco et al., 2019) and transported to the seminal vesicles via the vas deferens (Clements, 2000; Oliva et al., 2014; Rocco and Paluzzi, 2016). During mating, male *A. aegypti* deposit sperm from the seminal vesicles into the female reproductive tract initially in the bursa and is later transferred into the spermathecae for long-term storage (Camargo et al., 2020; Degner and Harrington, 2016). Herein, the results revealed lower spermatozoa counts in testes and seminal vesicles of males with *Itp* and *Itp-l* knockdown. Thus, this indicates that *Aedae*ITP/ITP-L knockdown reduces the incidence of blood feeding by females, and additionally, knockdown reduces the number of spermatozoa in male seminal vesicles and in the spermathecae of females, which ultimately results in fewer eggs laid and reduced larval hatching. It remains to be investigated exactly what role *Aedae*ITP and ITP-L play in male and female reproduction or in blood feeding behaviour. However, given the already established roles of these neuropeptides in *T. castaneum* reproduction (Begum et al., 2009), it is possible that *Aedae*ITP and ITP-L are similarly involved in regulation of male or female reproductive biology or mating behaviour, influencing successful mating and transfer of spermatozoa into the female *A. aegypti*. Indeed ITP and ITP-L are multifunctional neuropeptides involved in metabolism, regulation of water and ion homeostasis, cuticle expansion and melanization, and reproduction (Begum et al., 2009; Dircksen, 2009; Gáliková et al., 2018; Yu et al., 2016). The role in reproduction has been supported by ITPL expression in the *B. mori* male reproductive system with innervations of the accessory glands, the seminal vesicles, and the ejaculatory ducts (Klöcklerova, et al., 2023), all organs critical for successful mating and reproduction. ITPL expression has also been found in the seminal fluid of the brown plant hopper, *Nilaparvata lugens* (Yu et al., 2016), which is transferred to females during mating. Future research examining *Aedae*ITP and ITP-L signaling can provide greater insight on the actions of these neuropeptides in mosquito reproductive biology.

ITP and ITP-L peptides are highly homologous to the CHH peptides, which has been linked to molting, energy metabolism, immune defense, reproduction, and homeostatic regulation of osmotic and other stress responses (Sonobe et al., 2001; Webster et al., 2012). In locusts, ITP stimulates fluid reabsorption and Cl^-^, Na^+^, and K^+^ transport, while inhibiting secretion of H^+^ in the ileum (Audsley et al., 1992b; Meredith et al., 1996). In *Drosophila*, ITP plays an essential role in development and locomotion (Gáliková et al., 2018; Johard et al., 2009), and water homeostasis by protecting the fly from water loss by increasing thirst, reducing excretion rate, and promoting ingestion of water (Gáliková et al., 2018). Studies have also established functions of ITP during ecdysis in *M. sexta* (Drexler et al., 2007) and wing expansion in *N. lugens* (Yu et al., 2016), while ITP-L has been linked to ovarian maturation in *T. castaneum* (Begum et al., 2009) and produced as a seminal fluid protein in *N. lugens* (Yu et al., 2016). Differences in the primary structure and cellular distribution patterns of ITP and ITP-L peptides suggest they may serve different biological functions. In conclusion, the current results expand our understanding of ITP and ITP-L in insects, providing evidence of differential expression, insight into their cell-specific distribution, and revealing novel independent physiological roles for these neuropeptides in the *A. aegypti* mosquito. Importantly, these findings also contribute towards our understanding of *A. aegypti* reproductive biology, which is of medical importance given their propensity of feeding on human hosts and role as a vector of several viruses. As such, given these neuropeptides appear to hold pleiotropic actions related to successful mating and reproduction, further insights into ITP and ITP-L signaling could contribute towards development of novel strategies for decreasing the fitness of these vectors, that may improve control of these anthropophilic mosquitoes.

## Conflict of Interest

*The authors declare that the research was conducted in the absence of any commercial or financial relationships that could be construed as a potential conflict of interest*.

## Author Contributions

J.-P.P. and F.S. designed research; F.S. performed research; F.S. and J.-P.P. analyzed data; and F.S. and J.-P.P wrote and revised the manuscript.

## Funding

This research was funded by: Natural Sciences and Engineering Research Council of Canada (NSERC) Discovery Grant (JPP), Ontario Ministry of Research Innovation Early Researcher Award (JPP), and NSERC CGS-D (FS) and the Carswell Scholarships in the Faculty of Science, York University (FS).

## Acknowledgments

The authors sincerely thank Britney Picinic, Lulia Snan and Chiara Di Scipio for helping with mosquito collection and RNA extraction for expression profiles. We also sincerely thank Areej Al-Dailami, Profs. Ian Orchard and Angela Lange (University of Toronto Mississauga, Canada) for their generous training and support with confocal microscopy.

## Data Availability Statement

The datasets generated in this study can be found in the Borealis open data repository [https://borealisdata.ca/dataverse/PaluzziLab].

## Notes

### Competing Interest Statement

The authors have declared no competing interest.

