## Supplementary information for "Molecular characterization, localization, and physiological roles of ITP and ITP-L in the mosquito, *Aedes aegypti*"

**Table S1.** Gene-specific primer information used for amplification of *A. aegypti Itp* and *Itp-l* for RT-qPCR analysis, FISH probe synthesis and dsRNA synthesis. T7 promoter sequence is italicized.

| Oligo Name | Sequence (5' – 3') | Function | Product size |
|---|---|---|---|
| ***Itp* & *Itp-l* qPCR F** | AATCGCACTGGTACCTCTGG | qPCR amplification of *Itp* & *Itp-l* | 195bp (*Itp* & *Itp-l*) |
| ***Itp* qPCR R** | CCAAAACAGCCCTCTTTGCAG | qPCR amplification of *Itp* | |
| ***Itp-l* qPCR R** | GTGAAGCACGACTTTTTGCAG | qPCR amplification of *Itp-l* | |
| ***Itp* dsRNA F** | GACTGAATCAAATATGTGTTCCCG | Amplification of ds*Itp* target | 201bp |
| ***Itp* dsRNA R** | AACAACGAGTAGCAGTCTTCG | Amplification of ds*Itp* target | |
| ***Itp-l* dsRNA F** | CCAAACACGTTATAAGCAGCC | Amplification of ds*Itp-l* target | 486bp |
| ***Itp-l* dsRNA R** | ACGTCAAGATTTCCGGATGG | Amplification of ds*Itp-l* target | |
| ***Egfp* dsRNA F** | ACTCGTGACCACCCTGACCTACG | Amplification of ds*Egfp* target | 324bp |
| ***Egfp* dsRNA R** | AGATCTTGAAGTTCACCTTGATGCC | Amplification of ds*Egfp* target | |

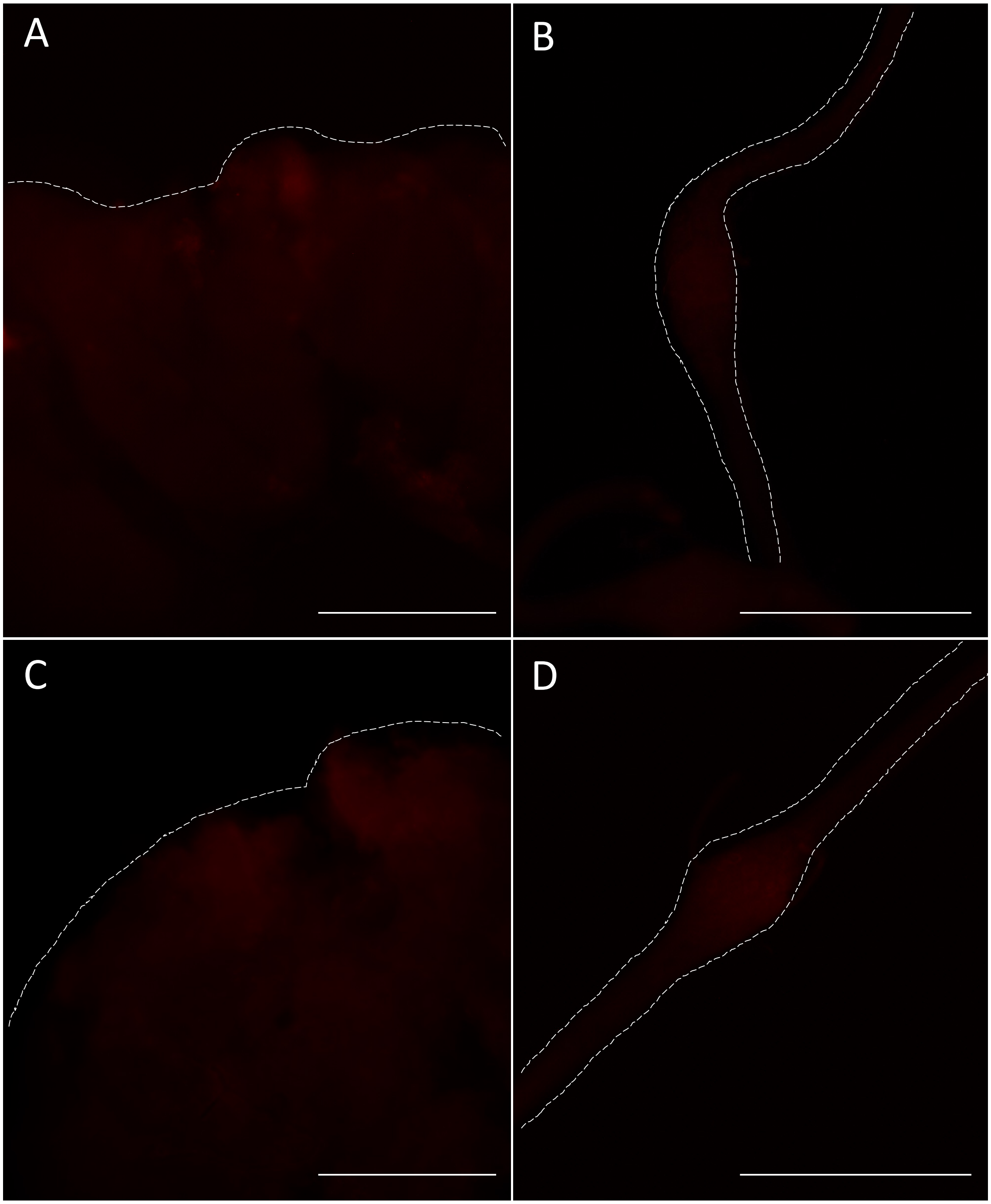


**Figure S1.** *Aedae*ITP and *Aedae*ITP-L preabsorbed controls and no primary controls. Central nervous system tissues were incubated in either (A,B) anti-*Aedae*ITP/ITP-L primary antiserum preincubated with 10 μM antigen or (C,D) no primary control. No staining was observed in the nervous tissues including the brain (A,C) or abdominal ganglia (B,D). Scale bars: (A,C) 200 μM; (B,D) 100 μM.

A

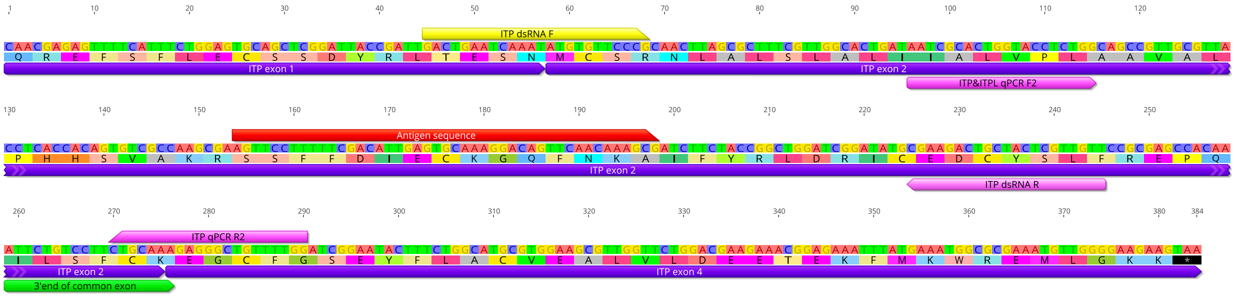


B

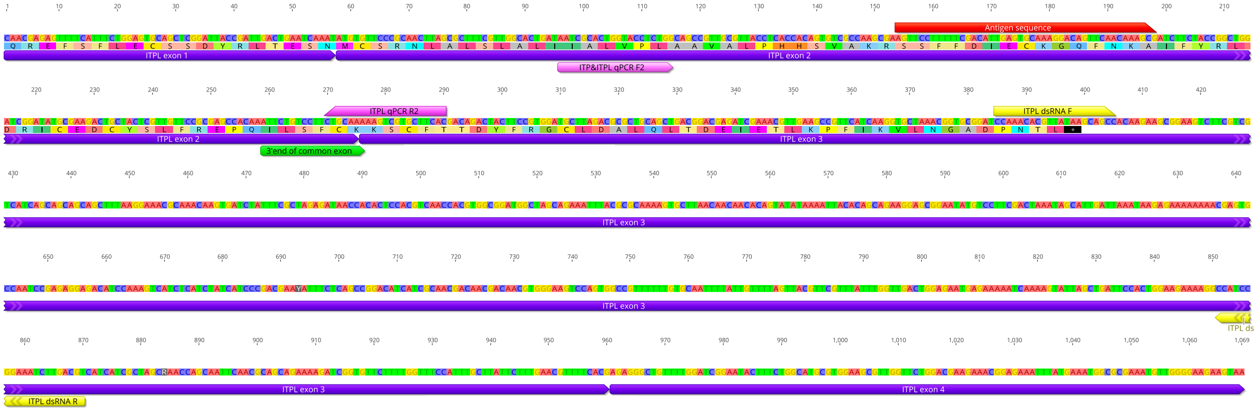


C

```
                      10         20         30         40         50         60         70         80
Consensus    SSFFDIECKG QFNKAIFYRL DRICEDCYSL FREPQILSFC KXXCFXXXYF XXCXXALXLX XEXEXXXXXX XXLXXXDPNT L  81
AedaeITP     SSFFDIECKG QFNKAIFYRL DRICEDCYSL FREPQILSFC KEGCFGSEYF LACVEALVLD EETEKFMKWR EMLGKK---- -  76
AedaeITP-L   SSFFDIECKG QFNKAIFYRL DRICEDCYSL FREPQILSFC KKSCFTTDYF RGCLDALQLT DEIETLKPFI KVLNGADPNT L  81
```

**Figure S2.** Primer sets mapped to *A. aegypti Itp* and *Itp-l* transcripts, and sequence alignment of *Aedae*ITP and *Aedae*ITP-L deduced peptides. (A) The *Itp* transcript coding sequence spans three exons, and (B) *Itp-l* spans four exons, denoted in purple. Targets for dsRNA synthesis included regions bracketed by primers shown in yellow and qPCR analysis shown in pink. Amino acid region encoding the antigen target sequence of custom *Aedae*ITP/ITP-L primary antisera is shown in red. The 3' end of common exon for *Itp* and *Itp-l* is shown in green. Top letters indicate nucleotides (positions of which are denoted by numbers above). Second panel of letter indicates the translated sequence. (C) Aligned amino acid sequences of ITP and ITP-L from *A. aegypti*, ITP (Genbank: AY950503), and ITP-L (Genbank: AY950506). Highlighting of residues indicates % identity with grey denoting 100% sequence identity.

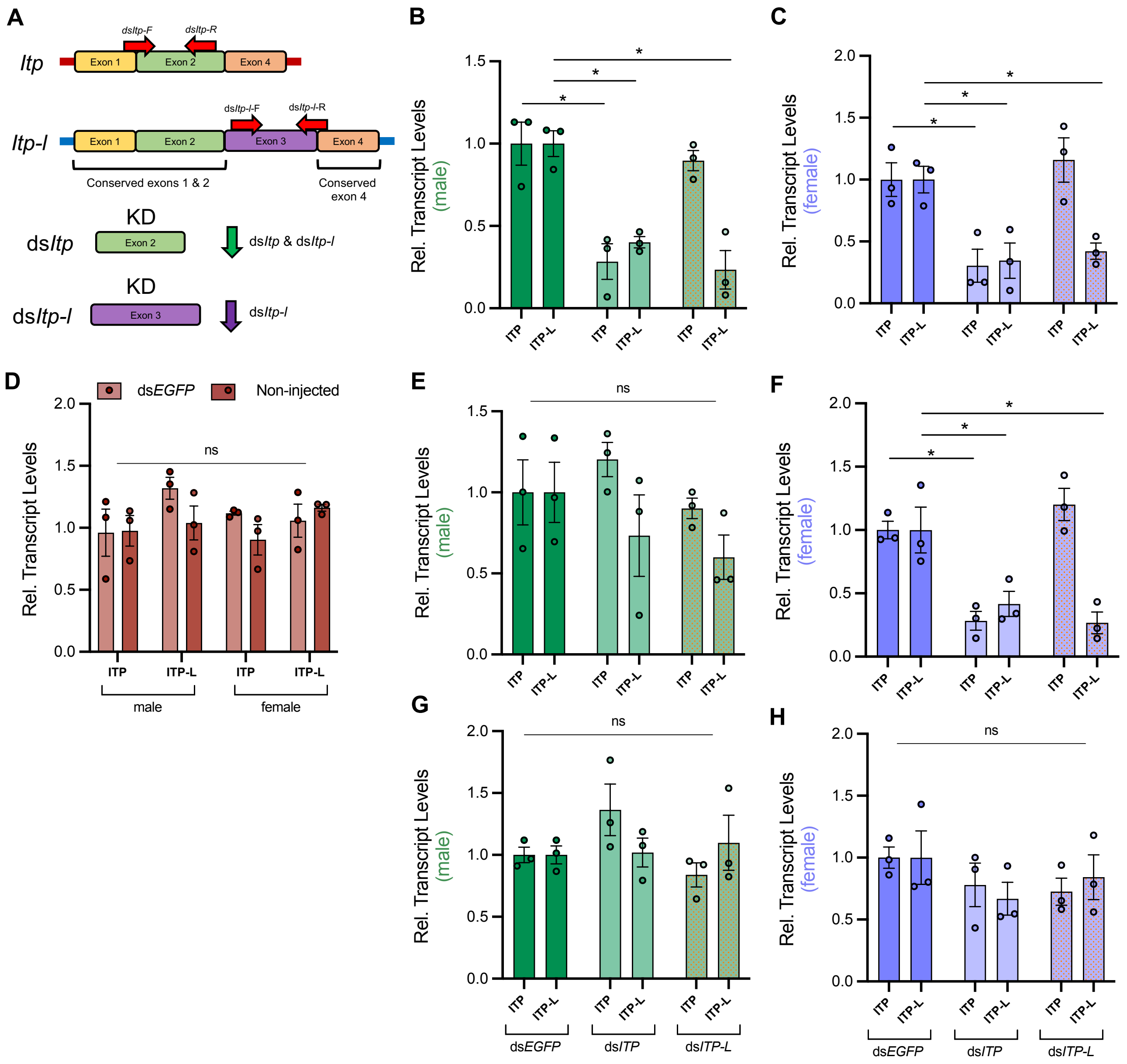


**Figure S3.** *AedaeItp* and *AedaeItp-l* knockdown efficiency in adult *A. aegypti*. (A) ds*AedaeItp* primers were designed over a common exon 2 resulting in the knockdown of both *AedaeItp* and *AedaeItp-l* transcripts, whereas ds*AedaeItp-l* primers were designed over a unique exon 3 targeting only the *AedaeItp-l* transcript. One day old adult male (B,E,I) and female (C,F,H) mosquitoes were injected with ds*AedaeItp*, ds*AedaeItp-l* or control ds*Egfp* (D) and collected four (B,C) six (E,F) and eight (G,H) days post-injection to determine knockdown efficiency. Transcript levels are shown relative to control ds*Egfp* mosquitoes. (D) ds*Egfp*-injected animals compared to non-injected animals. Data shown as mean±SEM; one-way ANOVA with Tukey's multiple comparison, asterisk (*) denotes significant knockdown, $p<0.05$, n=3 biological replicates (ns denotes no statistical significance).

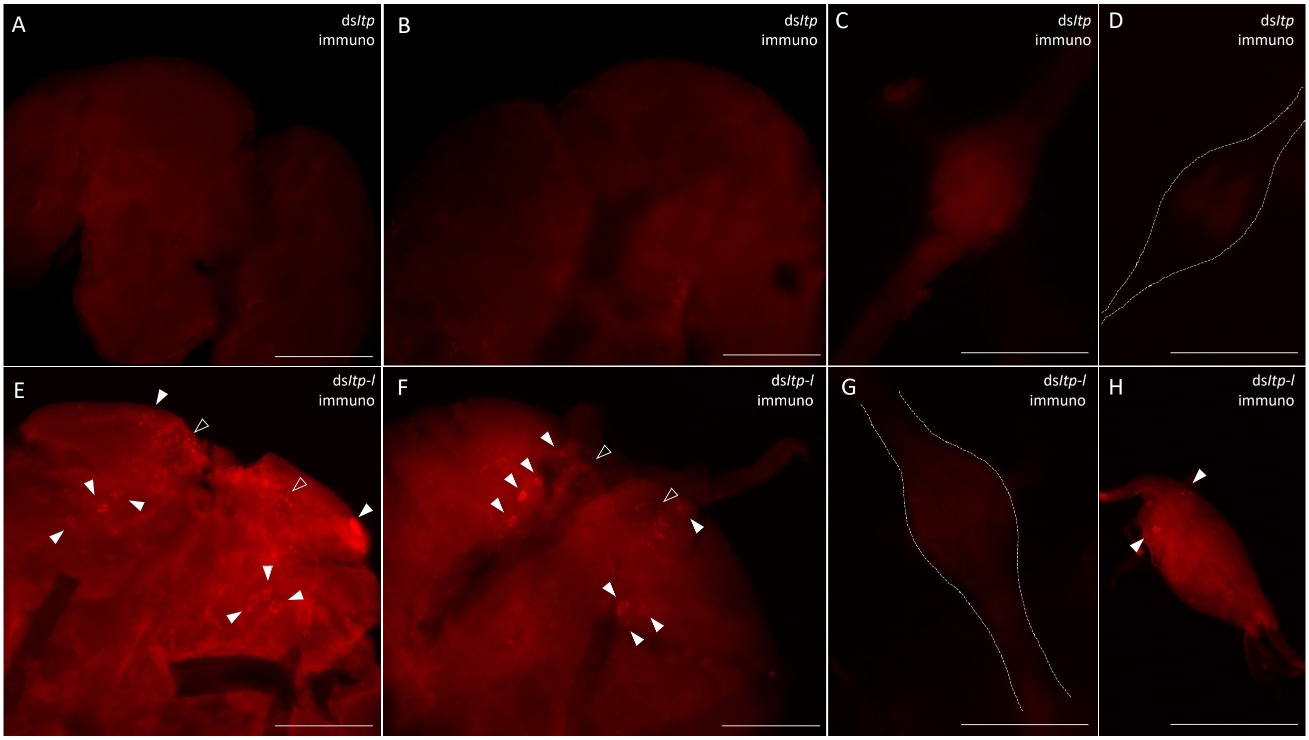


**Figure S4.** Immunolocalization of *Aedae*ITP and *Aedae*ITP-L in the central nervous system of adult *A. aegypti* mosquitoes following ds*AedaeItp* and ds*AedaeItp-l* injection. *Aedae*ITP- and *Aedae*ITP-L-like immunoreactivity is abolished in (A) ds*AedaeItp*-injected male and (B) female brain, (C) abdominal ganglia and (D) terminal ganglion. (E) ds*AedaeItp-l* treated male and (F) female brain reveal immunoreactivity in four pairs of neurosecretory cells (indicated by white arrowheads), with axonal processes emanating anteriorly towards release sites (indicated by empty arrowheads). (G) No immunostaining was observed in the abdominal ganglia following ds*AedaeItp-l* treatment whereas (H) one pair of lateral anterior cells were observed in the terminal ganglion. Scale bars: (A,B,E,F) 200 μM; (C,D,G,H) 100 μM.

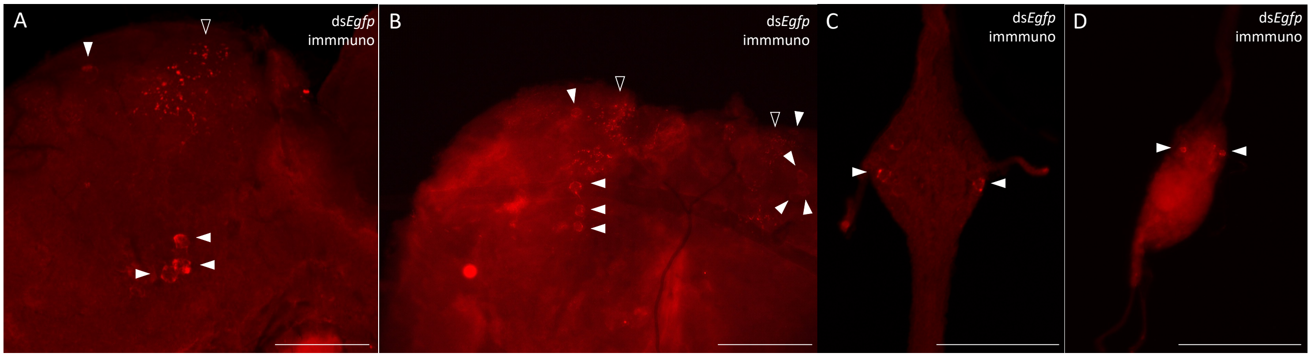


**Figure S5.** Immunolocalization and distribution of both *Aedae*ITP and *Aedae*ITP-L peptide in the central nervous system of adult ds*Egfp*-injected *A. aegypti* mosquitoes. ds*Egfp*-injected (A) male and (B) female mosquitoes reveals immunostaining in four pairs of neurosecretory cells (indicated by white arrowheads), with processes projecting anteriorly towards varicosities and blebs on the periphery of the brain (indicated by empty arrowheads). (C) Ventral view of abdominal ganglia and (D) terminal ganglion showing a single pair of lateral neurosecretory cells. Scale bars: (A,B) 200 μM; (C,D) 100 μM.

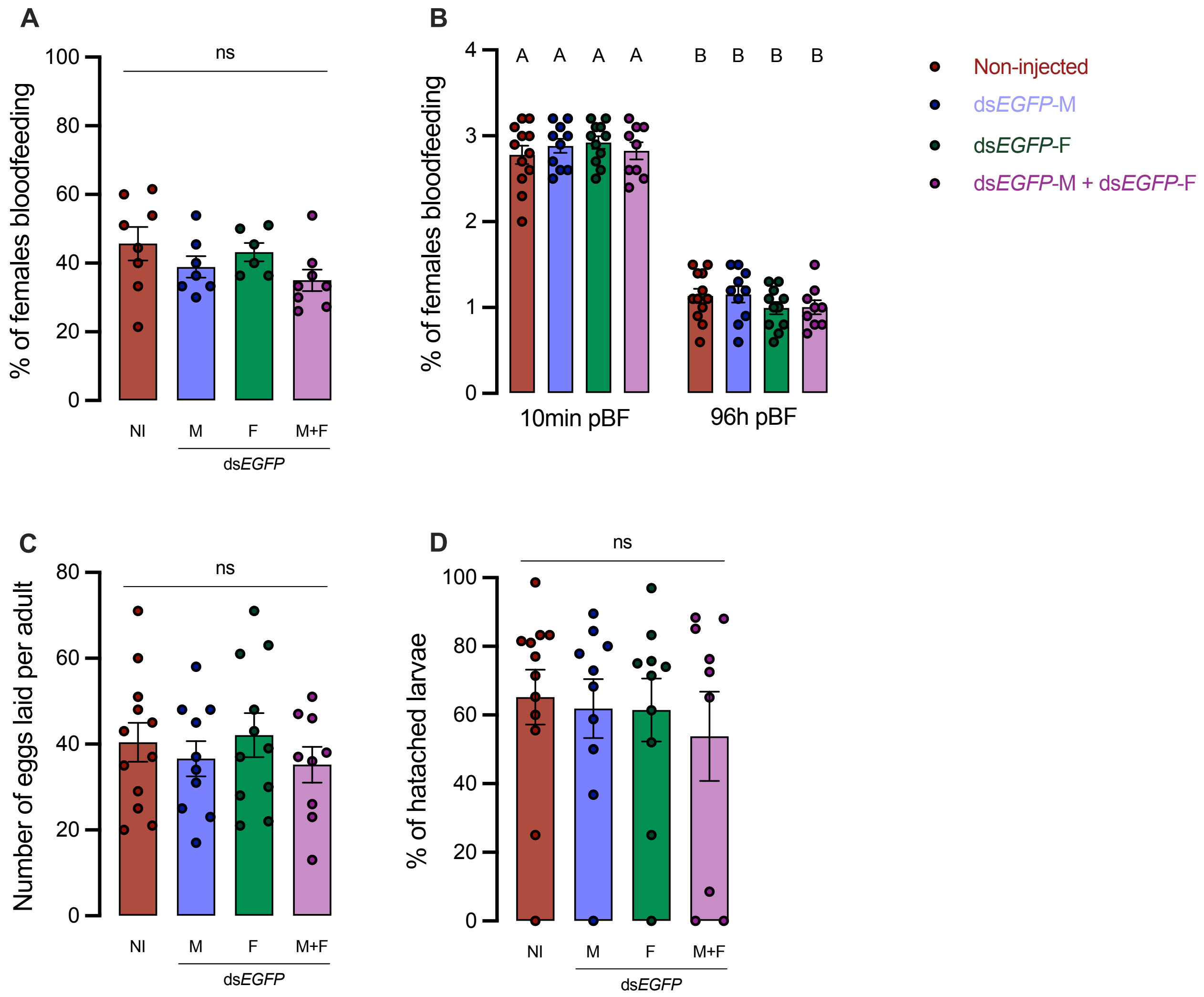


**Figure S6.** Effect of control ds*Egfp* on blood feeding, egg laying, and larval hatching (egg viability) in comparison to non-injected adult *A. aegypti*. The effect of control ds*Egfp* knockdown was tested on (A) preference for blood feeding, (B) weight of blood fed female before and after egg-laying, (C) number of eggs laid, and (D) percentage of larval hatching. Abbreviations: non-injected (NI), male (M) and female (F). Data labeled with different letters are significantly different from non-injected adults (mean±SEM; one-way ANOVA with Bonferroni multiple comparison, $p<0.05$, n=6–12 mating replicates, each point represents individual replicate values), (ns denotes no statistical significance).

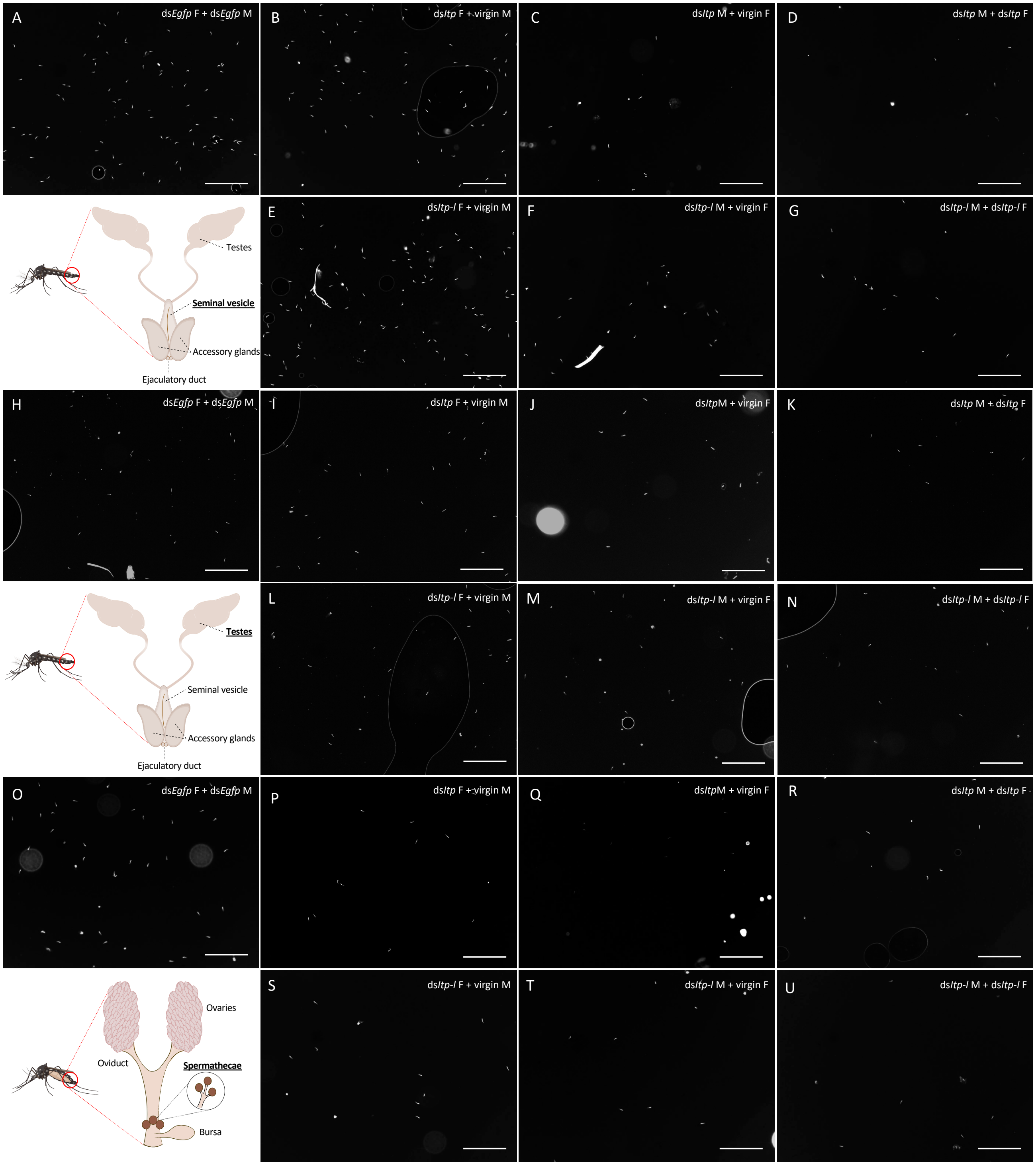


**Figure S7.** Representative images of immunofluorescent nuclei of total spermatozoa in the male seminal vesicle and testes, and female spermathecae of adult *A. aegypti* following RNAi (dsRNA)-mediated knockdown of *AedaeItp* or *AedaeItp-l*. Images of immunofluorescent nuclei (DAPI) from fixed spermatozoa in 1 µl droplet of DPBS with sperm from the seminal vesicle (A-G), testes (H-N) and spermatheca (O-U) of ds*AedaeItp* or ds*AedaeItp-l* injected animals from different mating treatments. Abbreviations: male (M) and female (F). Scale bar: 200 µm.

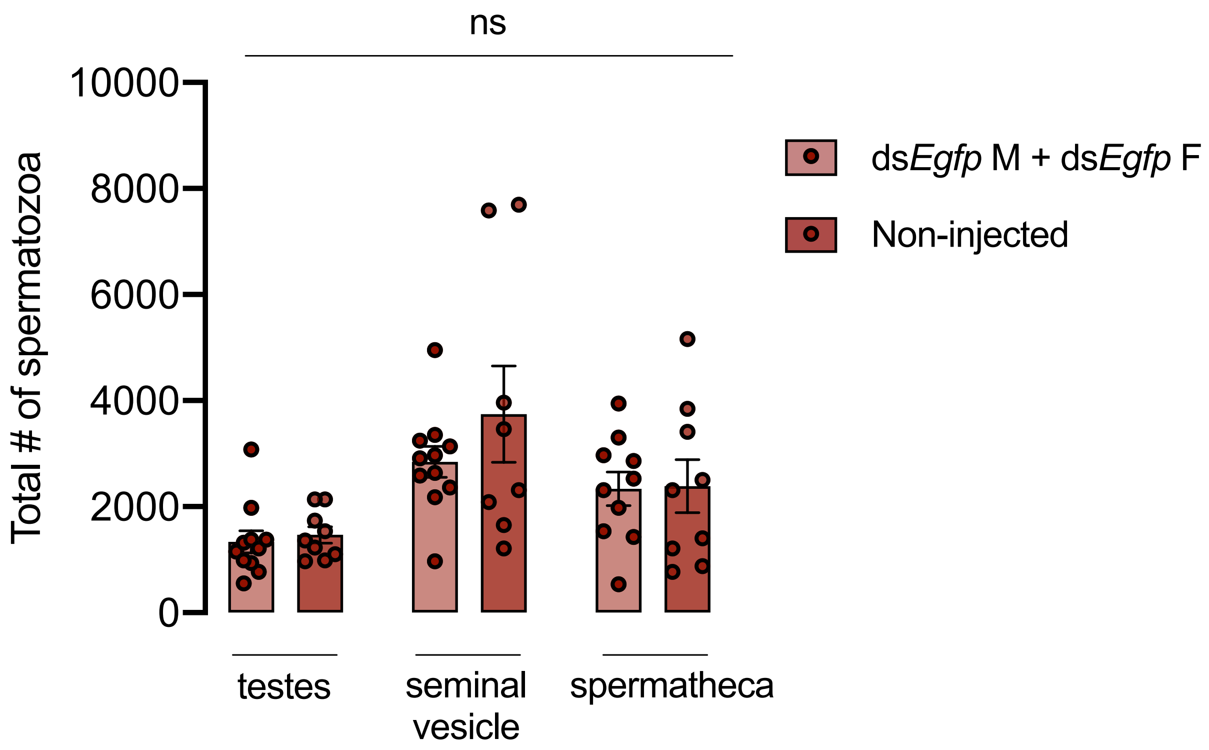


**Figure S8.** Total spermatozoa in the male testes and seminal vesicle and female spermathecae of adult *A. aegypti* following ds*Egfp* injection. Number of spermatozoa per adult four-days post ds*Egfp* injection and non-injected four-day old adults (mean± SEM; one-way ANOVA with Bonferroni multiple comparison, $p<0.05$, n=8–11 mating replicates, each point represents individual replicate values), (ns denotes no statistical significance).
